# High-frequency longitudinal white matter diffusion- & myelin-based MRI database: reliability and variability

**DOI:** 10.1101/2022.12.01.518514

**Authors:** Manon Edde, Guillaume Theaud, Matthieu Dumont, Antoine Théberge, Alex Valcourt-Caron, Guillaume Gilbert, Jean-Christophe Houde, Loika Maltais, François Rheault, Federico Spagnolo, Muhamed Barakovic, Stefano Magon, Maxime Descoteaux

## Abstract

Assessing the consistency of quantitative MRI measurements is critical for inclusion in longitudinal studies and clinical trials. Intraclass coefficient correlation and coefficient of variation were used to evaluate the different consistency aspects of diffusion- and myelin-based MRI measures. Multi-shell diffusion and inhomogeneous magnetization transfer datasets were collected from twenty healthy adults at a high-frequency of five MRI sessions. The consistency was evaluated across whole bundles and the track-profile along the bundles. The impact of the fiber populations on the consistency was also evaluated using the number of fiber orientations map. For whole and profile bundles, moderate to high reliability of diffusion and myelin measures were observed. We report higher reliability of measures for multiple fiber populations than single. The overall portrait of the most consistent measurements and bundles drawn from a wide range of MRI techniques presented here will be particularly useful for identifying reliable biomarkers capable of detecting, monitoring and predicting white matter changes in clinical applications and has the potential to inform patient-specific treatment strategies.

**Key points:**

- Reliability and variability are excellent to good for DWI measurements, and good to moderate for MT measures for whole bundles and along the bundles.
- The number of fiber populations affects the reliability and variability of the MRI measurements.
- The reliability and variability of MRI measurements are also bundle dependent.

## 1. Introduction

In recent decades, there has been a growing body of evidence that white matter (WM) plays a prominent role in various pathologies. In this context, longitudinal studies of WM have become increasingly important. Currently, various magnetic resonance imaging (MRI) techniques are used to assess different properties of WM tissues such as axonal density, fiber organization, or myelin (Jones et al., 2013). The most common technique is diffusion-weighted imaging (DWI), which includes single-compartment models such as Diffusion Tensor Imaging (DTI, Basser et al., 1994) or more complex models using multi-compartment fitting such as High Angular Resolution Imaging (HARDI, Dell’Acqua & Tournier, 2019; Jeurissen et al., 2013) and Neurite Orientation Density and Dispersion Imaging (NODDI, H. Zhang et al., 2012) from multi-shell diffusion MRI. Other techniques like magnetization transfer imaging (MTI, M. Kim & Cercignani, 2014; Wolff & Balaban, 1989) are also increasingly used to examine changes in the myelin content of WM. Key measures of WM microstructure derived from these models are sensitive to changes in a healthy population (Alexander, 2017; Beck et al., 2021; Boukadi et al., 2019; Honnedevasthana Arun et al., 2021; Koshiyama et al., 2020; Munsch et al., 2021; Uddin et al., 2019) or in pathological conditions (Beaudoin et al., 2021; Brown et al., 2017; Granziera et al., 2021; Laule & Moore, 2018; Lu et al., 2021; Rahmanzadeh et al., s. d.; Schneider et al., 2017). In addition, studies have also shown the effect of a treatment or therapeutic intervention on WM measures in clinical trials (Arnold et al., 2022; Gurevich et al., 2018; Roy et al., 2021; Vavasour et al., 2019).

However, reliably evaluating, monitoring, or predicting any changes in WM microstructure requires data with high consistency and enough statistical power to detect these changes (Poldrack et al., 2017; Zuo et al., 2019). MRI measurements can be influenced by random effects introducing measurement errors such as image noise, MRI signal variation or scanner model (Wang et al., 2012). In addition, repeated measure analyses also introduce additional sources of variation (a change of technologist during data acquisition or subject positioning, for example). Together, these errors affect data consistency which is an important factor in the sensitivity and specificity of the analysis (Tofts et al., 2018; Wang et al., 2012; Zuo et al., 2019). Therefore, it is essential to evaluate the different aspects of the consistency of the measurements including reliability, reproducibility and variability of measurements derived from MR images, especially so for the more novel and more complex quantitative MRI protocols. Here, reliability refers to the overall consistency of the measurements across subjects, i.e., it reflects both the degree of correlation and agreement between measures (Bruton et al., 2000; Koo & Li, 2016). Variability can be separated into within-subject variability and between-subject variability. Within-subject variability can be used to assess the ability to obtain similar values across sessions of the same subject, i.e., an index of measurement reproducibility. Finally, between-subject variability represents the sample heterogeneity, i.e., how much one subject differs from another.

To date, consistency of MRI-based WM measurements has been evaluated through numerous studies, especially for the DWI technique (Boekel et al., 2017; Grech-Sollars et al., 2015; Hakulinen et al., 2021; Magnotta et al., 2012; Teipel et al., 2011; Thieleking et al., 2021; Veenith et al., 2013). These studies reported moderate to high reliability in the WM using Intraclass correlation coefficient (ICC) or Pearson’s correlation ranging from 0.5 to >0.8 as well as within- and between-subject coefficients of variation (CV) ranging from 1 to 8% and 1 to 15% respectively. Among DTI-derived measures, Fractional anisotropy (FA) and Mean Diffusivity (MD) generally show the highest reliability across different WM regions (Acheson et al., 2017; Hakulinen et al., 2021; Luque Laguna et al., 2020; Palacios et al., 2017; Shahim et al., 2017; Thieleking et al., 2021; Zhou et al., 2018). For NODDI-derived measures, studies reported similar (intracellular volume fraction, ICvf) or higher (orientation dispersion, OD) reliability compared to DTI measures, while isotropic volume fraction (ISOvf) showed the poorest reliability (ICC<0.6) (Andica et al., 2020; Chung et al., 2016; Granberg et al., 2017; Lucignani et al., 2021; Tariq, 2013). In contrast, to the best of our knowledge, the reliability of HARDI-derived measurements such as apparent fiber density (AFD, D. Raffelt et al., 2012) and the number of fiber orientations (NuFO, Dell’Acqua et al., 2013a) has not been yet examined in healthy subjects or clinical population. Regarding MTI, MTR (Henkelman et al., 2001; Vavasour et al., 2011) - the most common measure - has been shown to have good reliability (ICC > 0.7) (Filippi et al., 2000; Hickman et al., 2004; Schwartz et al., 2019; van der Weijden et al., 2021; Weiskopf et al., 2013). More recently, Inhomogeneous magnetization transfer (ihMT, Varma et al., 2015) - a novel development of MT - has been shown to be more specific to myelin content compared to MTR (Duhamel et al., 2019; O. M. Girard et al., 2015; Manning et al., 2017; Varma et al., 2015; L. Zhang et al., 2020) and sensitive to multiple sclerosis-related (MS) processes in transversal studies (Obberghen et al., 2018; Rasoanandrianina et al., 2020; L. Zhang et al., 2020). To date, two studies reported good reliability of ihMT measuring with ICC ranging from 0.6 to 0.95 in different regions of the WM (Mchinda et al., 2018; L. Zhang et al., 2019), whereas the only longitudinal study suggests that ihMT may not have enough statistical power to detect changes during brain development (Geeraert et al., 2019). Hence, that reinforces the need to examine the reliability and variability of this recent, but promising, technique.

Nevertheless, several important issues that remain to be addressed : (1) although the recent studies include a reasonably large sample size n ≥ 20 (Boekel et al., 2017; Hakulinen et al., 2021; Lehmann et al., 2021; Thieleking et al., 2021), most of them are based on limited data with sample sizes of n ≥ 10 (Andica et al., 2020; Chung et al., 2016; Granberg et al., 2017; Koller et al., 2020; Tariq, 2013; L. Zhang et al., 2019); (2) most of the previous studies are focused on a short-period (scan-rescan within a week or with 2-4 weeks intervals) rather than longer time intervals. Indeed, longitudinal neuroimaging studies or clinical trials are typically separated by a few weeks (> 3 weeks) to several months; (3) only a few studies include more than two or three-time points (Cai et al., 2021; Schwartz et al., 2019), thus not generating enough data per subject to assess relevant reliabilities; and (4) few studies have directly compared multiple WM microstructural from several MRI techniques in the same population (Koller et al., 2020; Schwartz et al., 2019).

On the other hand, none of these studies evaluated the impact of local WM complexity on consistency, particularly the number of fiber populations. Indeed, these studies consider each voxel as a single entity with a homogeneous fiber population. However, it has been described that voxels contain multiple fiber populations *i*.*e*., between 66% to 90% of white matter voxels cannot be assumed to contain a single coherently oriented axon bundle (Jeurissen et al., 2013; Volz et al., 2018). In addition, Volz and colleagues have recently shown that the value of FA depends on the number of fibers considered in the voxel, with a greater FA value for the unidirectional fiber population and smaller when the multidirectional fiber population is considered (Volz et al., 2018). Thus, measurements derived from different models - and by extension, their consistencies - may vary depending on the underlying WM organization, especially within bundle or track-profiles.

To address these problems, we designed a repeated-measure study to collect multiple microstructural (anatomical, multi-shell diffusion and inhomogeneous MT) MRI datasets in “high-frequency” - *i*.*e*., a high number of MRI acquisitions over a short period of time (six months) for twenty healthy subjects. All subjects were scanned five times with an average interval of four weeks for a total of 100 MRIs. This high-frequency dataset thus generates enough data per subject to optimize a relevant assessment of the consistency of different brain MRI measurements. The reliability and variability were evaluated using the Intraclass Coefficient correlation (ICC) value and within- and between-subject coefficient of variation (CVw and CVb, respectively). Then, the consistency of MRI measurements was evaluated across the bundles as a tracts-of-interest analysis approach - *i*.*e*., averaging voxels within each WM bundle. To go further, the consistency of each WM measure was also evaluated as a profile along the bundle using *tractometry* (Cousineau et al., 2017; Yeatman et al., 2012). Finally, the same analyses were carried out by splitting each white matter bundle mask according to the number of fiber orientations using the NuFO map, a useful index of the number of fiber populations.

## 2. Methods

Method and results are documented and available at https://high-frequency-mri-databasesupplementary.rtfd.io.

### 2.1. Participants

Twenty healthy adults (mean age 36 years, age range 29-46 (SD = 4.7), 3 men and 17 women) were recruited from the environment of the University of Sherbrooke and the Centre Hospitalier Universitaire of Sherbrooke (CHUS). The study was designed with this proportion of male and female subjects to match a future MS group. Participants were screened for eligibility to undergo MRI, no history of brain disease or injury, left-handedness and received financial compensation for their participation. The study was approved by the ethics committee of the CHUS (Comité d’éthique de la recherche du CIUSSS de l’Estrie) in Sherbrooke, Canada and all participants gave prior informed written consent.

### 2.2. MRI data acquisition

Whole-brain MRI data were acquired using a clinical 3T MRI scanner (Ingenia, Philips Healthcare, Best, Netherlands) using a 32-channel head coil. Each MRI session lasted approximately 33 min and was repeated 5 times over 6 months and a 4-week interval (+/-1 week). For each participant, images were acquired at approximately the same time of day to avoid potential diurnal effects (*i*.*e*., a morning participant had all sessions in the morning, with a tolerated 2–3-hour variation). All MRI data acquisitions were aligned on the anterior commissure-posterior commissure plan (AC-PC) and included (a) anatomical 3D T1-weighted, (b) multi-shell diffusion-weighted images (DWI), (c) inhomogeneous magnetization transfer (ihMT).

a. 3D T1-weighted MPRAGE image was acquired axially at 1.0 mm isotropic resolution, repetition time (TR)/echo time (TE)/inversion time (TI) = 7.9/3.5/950 ms, field-of-view (FOV) = 224×224 mm^2^ yielding 150 slices, flip angle = 8° for an acquisition time of 4 min 20 s.
b. Multi-shell DWI images were acquired with a single-shot EPI spin-echo sequence at 2.0 mm isotropic resolution, TR/ TE= 4800/92 ms, SENSE factor = 1.9, Multiband-SENSE factor = 2, flip angle of 90°, FOV=224×224 mm^2^, 66 slices for an acquisition time of 9 min 19s. The data comprised of 100 unique directions uniformly spread over three shells at b = 300 mm^2^/s (n=8 directions), b = 1000 mm^2^/s (n=32 directions), b = 2000 mm^2^/s (n=60 directions), with non-diffusion-weighted images b = 0 mm^2^/s (n=7), for a total of 107 total diffusion volume (Caruyer et al., 2013). To correct EPI distortions, a reversed phase-encoded b = 0 image was acquired right after the DWI acquisition, with the same geometry (Andersson et al., 2003).
c. Inhnomogeneous MT images were acquired using a 3D segmented-EPI gradient-echo sequence with different MT preparation pulses with first TR/TE = 3.6/112 ms, 2 × 2 mm resolution, flip angle of 15°, FOV=224×224 mm, 65 slices of 2 mm of thickness and 3 echoes with echo spacing 6.0 ms for an acquisition time of 6 min 04s.Inhomogeneous MT uses a magnetization preparation (10 Hann pulses of 0.9 ms duration with 1.5 ms interval at a frequency offset of +/-7000 Hz) (Varma et al., 2015). Two additional reference images were acquired for each echo without MT preparation, with the same parameters as the MT sequence and a second with a higher flip angle (30°) and a shorter TR (20 ms) for quantification purposes.

### 2.3. MRI processing

#### 2.3.1. Tractoflow: DWI and T1 processing

After visual quality assessment, for each participant, *Tractoflow* (Theaud et al., 2020) a pipeline developed by SCIL (https://github.com/scilus/tractoflow), was used to process DWI and T1w images. This pipeline generates both diffusion measures and tractography of WM, from raw DWI, T1w, bvec/bval files and the reversed phase-encoded b=0 and has been proved to be highly reproducible in time and immediate test-retest (Theaud et al., 2020). Briefly, after denoising and correcting the raw DWI images for motion, eddy-currents, geometric distortions and field inhomogeneity, the fiber orientation distribution function (fODF) was generated using constrained spherical deconvolution (Descoteaux et al., 2007; Tournier et al., 2007) with a fixed fiber response of [15, 4, 4] × 10^−4^ s/mm^2^ for all subjects (Pierpaoli & Basser, 1996), as recommended in (Dell’Acqua et al., 2013b), a maximal spherical harmonics order of 8 and all b-value DWI data. Four DTI measures including Fractional Anisotropy (FA), Mean Diffusivity (MD), Radial Diffusivity (RD) and Axial Diffusivity (AD) were computed and HARDI-derived measures including total apparent fiber density (AFD total) and the number of fiber orientations (NuFO), were extracted from the fODF (see Table 1 for a list of available measures). In parallel, the T1w anatomical image was also denoised, corrected, and registered to the b=0 and the FA images before tissue segmentation to generate the tracking maps including inclusion, exclusion maps and a WM seeding mask (G. Girard et al., 2014). The whole-brain ensemble tractogram was generated from an fODF map and tracking masks using both the anatomically constrained particle filter tracking algorithm (PFT, G. Girard et al., 2014) and local tracking with 5 and 2 seeds per voxel respectively. Except for the number of seeds per voxel, all parameters used the default *Tractoflow* settings (see Theaud et al., 2020 for a complete pipeline description).

**Table 1.** List of pipelines, models and measures evaluated with the corresponding abbreviations.

| Public pipeline | Models | Measures | Abbreviation |
| --- | --- | --- | --- |
| Tractoflow<br>https://github.com/scilus/tractoflow | Diffusion Tensor Imaging (DTI) | Fractional Anisotropy | FA |
|  |  | Axial Diffusivity | AD |
|  |  | Radial Diffusivity | RD |
|  |  | Mean Diffusivity | MD |
|  | High Angular Resolution Diffusion Imaging (HARDI) | Apparent fiber density total | AFD total |
|  |  | Number of Fiber Orientation | NuFO |
| NODDI Flow<br>https://github.com/scilus/noddi_flow | Neurite Orientation Dispersion and Density Imaging (NODDI) | Intracellular Volume Fraction | ICVF |
|  |  | Extracellular Volume Fraction | ECVF |
|  |  | Isotropic Volume Fraction | ISOVF |
|  |  | Orientation Distribution | OD |
| ihMT flow<br>https://github.com/scilus/ihmt_flow | Magnetisation Transfer Imaging (MTI) | inhomogeneous Magnetization Transfer (MT) ratio | ihMTR |
|  |  | inhomogeneous MT delta R1 saturation | ihMTdR1sat |
|  |  | Magnetization Transfer ratio | MTR |
|  |  | Magnetization Transfer saturation | MTsat |

#### 2.3.2. Neurite Orientation Dispersion Density Imaging

NODDI is a multi-shell compartment modelling technique that identifies three types of microstructural environments: intracellular, extracellular, and CSF compartments (Zhang et al. 2012). NODDI measures were extracted using *NODDI flow* from SCIL (https://github.com/scilus/noddi_flow), which used Accelerated Microstructure Imaging via Convex Optimization (AMICO, Daducci et al., 2015) from multi-shell DWI images. Complementing, the extracellular volume fraction (ECvf), a measure of the volume fraction within a voxel that is not neuronal and assumed to be due to glial cells infiltration, was computed as follows ECvf=1-ICvf. Finally, four microstructural maps were generated: ECvf, intracellular volume fraction (ICvf), isotropic volume fraction (ISOvf), and orientation dispersion (OD).

#### 2.3.3. Magnetization Transfer Imaging (MTI)

Inhomogeneous MT images were processed using a custom in-house pipeline including tools from the FSL, Advanced Normalization Tools software (ANTs, Avants et al., 2011) and SCIL pipeline scripts (https://github.com/scilus/ihmt_flow). For each echo, raw ihMT images were firstly co-registered using ANTs linear registration (Avants et al., 2008). Next, the reference image was used to perform tissue segmentation with the *AtroposN4* command from ANTs. Tissue maps from the above T1w processing were concatenated and used as brain mask during ihMT processing. Two ihMT images were generated from all frequencies and reference images as previously described in Varma et al., 2015: ihMT ratio (ihMTR) and ihMT ΔR1 saturation (ihMTdR1sat) – developed to enhance ihMTR contrast by decoupling the ihMTR signal from the T1 longitudinal relaxation rate. Additionally, from the positive frequency data and reference images, we also computed two “standard” MT images: MT ratio (MTR) and MT saturation (MTsat) – generated as described in Helm et al., 2008. However, it should be noted that these MT measures are computed from ihMT acquisitions whose saturation frequency of 7000 Hz is outside the range generally used for MTR and MTsat (1000-2500 Hz). Finally, the four resulting myelin-sensitive maps were registered to the b=0 and the FA images using nonlinear SyN ANTs. Table 1 provides the complete list of measures included in the analyses.

### 2.4. White matter virtual dissection

The major fascicles were automatically extracted using RecoBundlesX (Rheault, 2020) (https://zenodo.org/record/4104300#.YNoPlXVKiiM) a multi-atlas and multi-parameter version of RecoBundles (Garyfallidis et al., 2018). For the sake of clarity and to avoid overloading, we focus the paper on a subset of bundles to show consistency in one association, commissural and projection pathway: Arcuate Fasciculus (AF), section 3 of the Corpus Callosum (CC), and Cortico-Spinal Tract (CST) are selected as bundles of interest (Catani & Schotten, 2012). In addition, the inferior fronto-occipital fasciculus (IFOF) is also included to show consistency in a “hard-to-track” long bundle (Figure 1). Bundle colors will be matched throughout the results. All analyses are nonetheless conducted on all bundles and measures, and the respective results are available at https://high-frequency-mri-database-supplementary.rtfd.io/ and included m the supplementary data.

**Figure 1.**
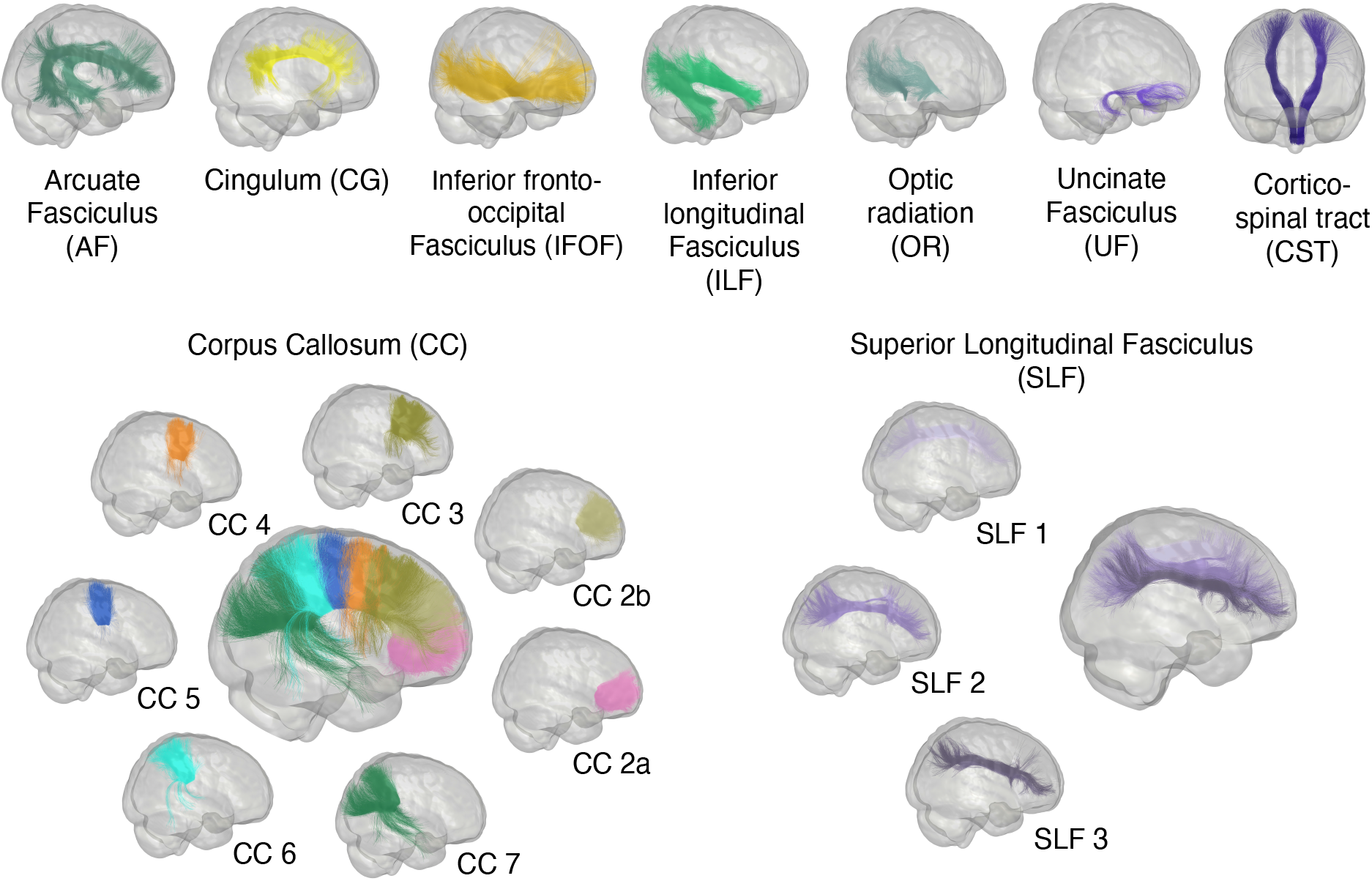
Representation of the major bundle models used by RecobundlesX as shape priors to extract the bundles from the whole tractogram. Bundles of both hemispheres are shown.

### 2.5. Common space and average measures

To perform consistency voxel-based analysis, all images were registered in a common space. Symmetric diffeomorphic normalization (SyN) of ANTs is used to build a template in diffusion space based on our population. For the registration, we used iterative rigid, affine, and SyN (neighborhood cross-correlation) transformations with optimal similarity measures for the linear (mutual information) (Avants et al., 2011) (https://high-frequencymri-database-supplementary.readthedocs.io/en/latest/pipeline/common_space.html). The processed b0 images resampled at 1 mm isotropic from *Tractoflow* were used as an input with four iterations with decreasing degrees of downsampling and smoothing.

All subject-specific measures maps and bundles were aligned in the common diffusion space using the resulting nonlinear registration. Finally, averaged maps were computed for each measure and shown in Figure 2 and available at https://high-frequency-mri-database-supplementary.readthedocs.io/en/latest/results/average_maps.html.

**Figure 2.**
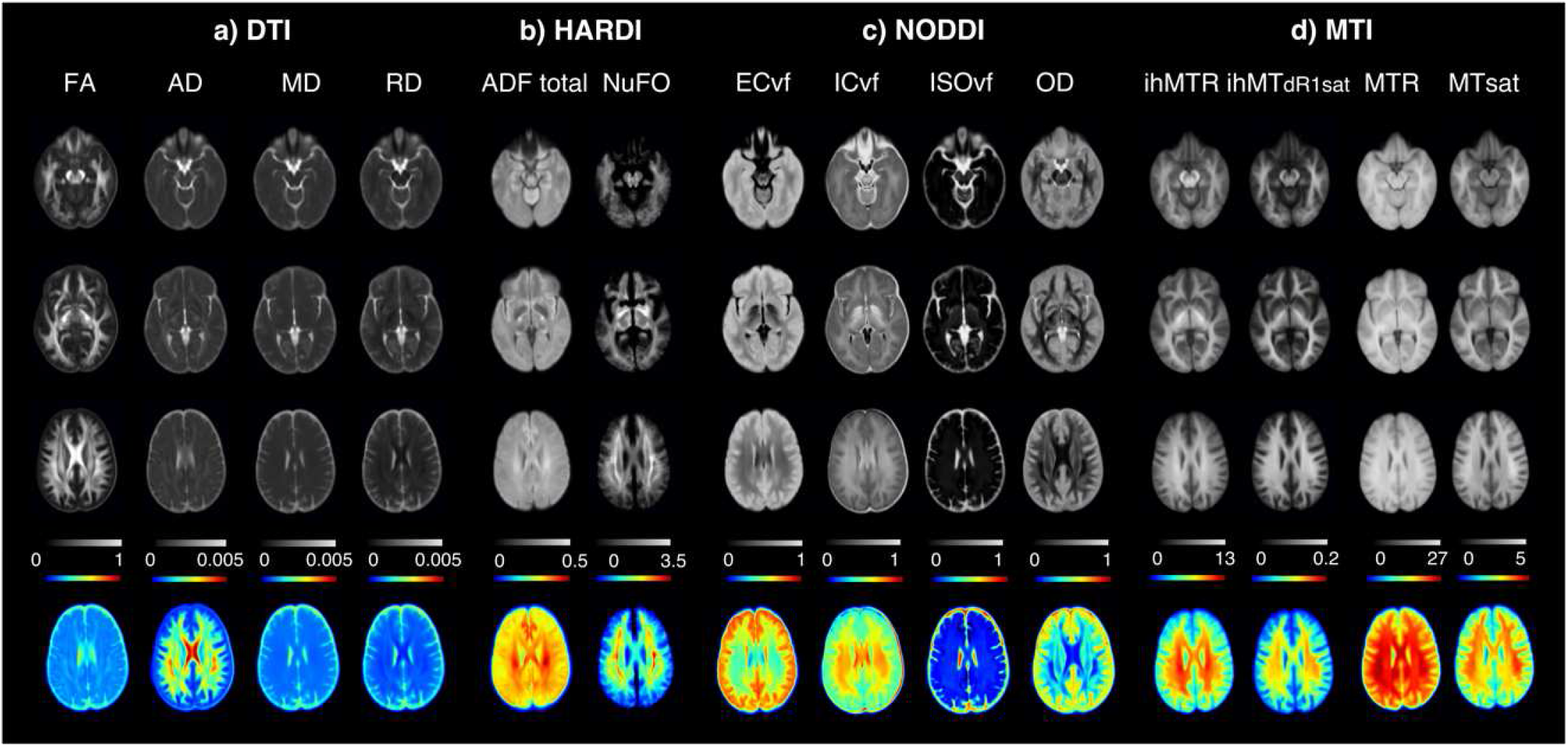
Microstructural maps, in a) DTI measures including FA (fractional anisotropy), AD (axial diffusivity, units = mm2/s), MD (mean diffusivity, units = mm2/s) and RD (radial diffusivity, units = mm2/s); b) HARDI measures with AFD total (total apparent fiber density) and NuFO (number of fiber orientations); c) NODDI measures including ECvf (extracellular volume fraction) and ICvf (intracellular volume fraction) compartment, ISOvf (isotropic volume fraction) and OD (orientation distribution) and d) MTI measures including ihMTR (inhomogeneous MT ratio), ihMTdR1sat (inhomogeneous MT saturation), MTR (MT ratio), MTsat (MT saturation); averaged across participant. All contrasts are registered to diffusion space.

### 2.6. Consistency for whole-bundle average and profiles along bundles

For each bundle, the consistency of the different measurements was evaluated from (1) the bundle-averaged *i*.*e*., one measure for the whole bundle and (2) along the bundle as a profile, also called track-profile or connectometry (Cousineau et al., 2017; Yeatman et al., 2012; Yeh et al., 2016). For the bundle-averaged, the density map was used to generate a binary mask of each whole bundle in the common space. Then, to minimize the effect of partial volume, each whole bundle mask was eroded by one voxel to generate a conservative bundle mask that we called the “safe mask”. For the consistency profiles, *Tractometry_flow*, a public pipeline developed by SCIL (https://github.com/scilus/tractometry_flow, Cousineau et al., 2017; Yeatman et al., 2012) was applied to each subject-specific bundle to obtain binary mask corresponding to each bundle section (each section corresponding to a specific label). Each bundle was resampled into 10 equidistant sections and intersected with the safe mask.

Next, left and right masks were merged for each average and section bundle mask (Supplementary Figure 1). Finally, DTI, HARDI, NODDI and MTI measures were extracted for each bundle mask over session.

### 2.7. Impact of fiber populations on consistency

Many studies have reported that voxels containing multiple fiber populations (Jeurissen et al., 2013; Volz et al., 2018) affect the microstructural measures (Volz et al., 2018). Here, we evaluate the effect of fiber populations projecting in multiple orientations on the consistency of the WM measures. To this end, we used the NuFO maps which are estimated from the number of local maxima of the fODF profile in each voxel (Dell’Acqua et al., 2013b). The intensity of each voxel corresponds to the number of fiber populations, ranging from 1 for the single fiber population to 2 and more for the multiple fiber populations (Jeurissen et al., 2013). We apply two thresholds of 1 and ≥2 on the NuFO map to compartmentalize the “average” bundle (i.e., whole bundle) into “single” and “multi” fiber populations compartments, respectively. For this, each voxel of the whole and section masks for each bundle was sorted according to these two thresholds (https://high-frequency-mri-database-supplementary.readthedocs.io/en/latest/pipeline/fiber_population.html.)

An overview of our analysis pipeline is illustrated in Supplementary Figure 2 and https://high-frequency-mri-database-supplementary.readthedocs.io/en/latest/data/overview.html#pipeline-summary.

### 2.8. Quality control

A visual quality assessment procedure was carried out for major steps including raw input data, preprocessing, registration steps, bundles segmentation and tract profiles using *dMRIqc flow* (https://github.com/scilus/dmriqc_flow). Because resampling the bundles involves smaller mask volumes and therefore, introduces a potential confounding factor, it is important to ensure that each section of bundles contains enough voxels to assess consistency measurements. For this purpose, we extracted the volume of each section corresponding to the bundle profile analyses and the fiber population compartments that generate a novel subdivision of the masks of each section in single- and multicompartment. Two minimum thresholds of 1000 and 400 voxels respectively were used to perform analyses, therefore, sections of bundles that had fewer voxels than these thresholds were excluded.

Since ISOvf accounts for the isotropic volume fraction, this parameter has generally very low values in the WM (Tariq, 2018), and many studies emphasized the poor reliability of this NODDI parameter (Andica et al., 2020; Chung et al., 2016; Lehmann et al., 2021; Lucignani et al., 2021). To improve the consistency of this parameter, we proposed an evaluation of different thresholds to remove values close to zero. A range of thresholds between 0 and 0.1 with a step size of 0.01 was used.

### 2.9. Evaluation of bundles

The evaluation of the reproducibility of bundles is carried out first to minimize the impact of the variability of reconstruction by tractography on the measurements. The reproducibility of the bundles was achieved using the same method as Rheault et al., 2020, 2022. We computed the Dice similarity score, correlation between the density maps and adjacency streamlines from all pairwise combinations to provide the agreement between segmentations of the same bundle across sessions.

### 2.10. Correlation analysis

Pearson’s correlation coefficient (r) was used to evaluate the covariance of the averaged diffusion measures for all bundles. For this, we used each bundle’s averaged measure, extracted from each voxel and averaged along bundles. Pearson correlations were computed for each session and then averaged across sessions to generate an average correlation map for all sessions. The average correlation interactive map and correlation interactive maps corresponding to the sessions are available at https://high-frequency-mri-database-supplementary.readthedocs.io/en/latest/results/correlation.html.

### 2.11. Consistency evaluation

#### 2.11.1. Consistency measures

The reliability of computed measures was investigated using the Image Intra-Class Correlation coefficient (I2C2, Shou et al., 2013), a generalization of the Intra-Class Correlation coefficient (ICC, Koo & Li, 2016) to n-dimensional images (one-way random effect, absolute agreement). The ICC estimates the correlation between measures values corresponding to different sessions in terms of their consistency across subjects. The variability induced by within-subject and between-subject effects on the measures was quantified using two coefficients of variation per measure. The coefficient of variation within-subject (CVw) was used to evaluate the dispersion of observations when repeatedly measuring a single individual (i.e., reproducibility), thus representing the amount of random error or noise contributing to the measure. For the CVw, the CV is first estimated per subject over their respective imaging sessions and then averaged. The coefficient of variation between-subject (CVb) was used to evaluate the sample heterogeneity. The CVb is obtained by first averaging each subject session-wise, to then estimate the CV over those averages.

Confidence intervals and p-values were obtained for the I2C2 using non-parametric bootstrap (Briggs et al., 1997; Efron & Tibshirani, 1994) and the accelerated bias-corrected percentile method (Diciccio & Romano, 1988; Efron & Tibshirani, 1994) using SciPy tools (https://scipy.org/). To avoid overloading, confidence intervals and p-values are only reported for selected bundles and the whole bundle analysis (see Supplementary Table 2).

#### 2.11.2. Voxel-based consistency analysis

Computation of consistency measures was restricted to the safe white matter masks. The consistency analyses of each measure were carried out at the voxel-level within the bundles mask. The individual masks corresponding to each subject and session in the common space were provided as input. The overlap between masks across sessions and subjects was then performed as described in the consistency measures. Incomplete overlap of mask between subjects and sessions was compensated by densifying each measure in the affected regions voxel-wise, using the average value estimated from the available subjects or sessions. The averaged masks used for the computation of statistical measurements are then obtained subject-wise or session-wise by mathematical union.

## 3. Results

### 3.1. Quality control

None of the data were excluded based on input quality controls. However, two subjects were excluded from IFOF and UF analyses due to failed reconstruction of these bundles - caused by poor WM-GM-CSF segmentation in the internal capsule. In this case, all sessions were excluded for these two bundles but included for the correctly reconstructed bundles. CC Part 1 reconstruction failed for most subjects and was therefore excluded from all analyses. The set of bundles finally included in this paper is shown in Figure 1. Regarding, the number of voxels in each section of bundles, section 10 of the cingulum bundle had an average volume under the threshold. The consistency profile of this bundle was therefore generated for sections 1 to 9 (Supplementary Figure 3). No other bundle sections were excluded based on volume. Finally, based on the graph and consistency results for the different thresholds of the ISOvf map, before analysis, an additional thresholding step was applied to exclude values under 0.045, for each subject (Supplementary Figure 4).

### 3.2. Diffusion- and myelin-based maps in diffusion template space

Three representative axial slices of the resulting averaged maps in common diffusion space are shown in Figure 2, the bottom row represents the third line with a colormap. As expected, the maps highlight the regional variation of measurements corresponding to each map, both in grey (top rows) and with a range of colors (bottom row). The maps are smooth but show sharp contrasts between different tissue types such as CSF, GM and WM. Moreover, higher values are observed in regions of highly structured white matter as can be seen with FA, AFD total, NuFO, ICvf or OD in corona radiata regions, internal capsule, forceps, or thalamic radiation, and inversely for ISOvf or MD. As for the other maps, higher ihMTR or ihMTdR1sat values are observed in these same regions. One can appreciate qualitatively the similarities between all the different diffusion and myelin-based measures. The resulting averaged maps in common space are also available at https://high-frequencymri-database-supplementary.readthedocs.io/en/latest/results/average_maps.html.

### 3.3. Consistency of bundle tractography

Dice scores presented in Figure 3 reveal that most of the bundles are highly reproducible (mean Dice > 0.7). Dice scores of voxels are high and close for all bundles, ranging from 0.56 (for SLF 1) to 0.84 (for CC 2a), which indicates that the overall spatial agreement is good. Across all bundles, SLF 1 exhibits the lowest Dice scores on average (density correlation and adjacency results are available at https://high-frequency-mridatabase-supplementary.readthedocs.io/en/latest/results/bundles_reproductibility.html). SLF 1 was not excluded from the analysis, but its results should be taken with caution (Figure 3).

**Figure 3.**
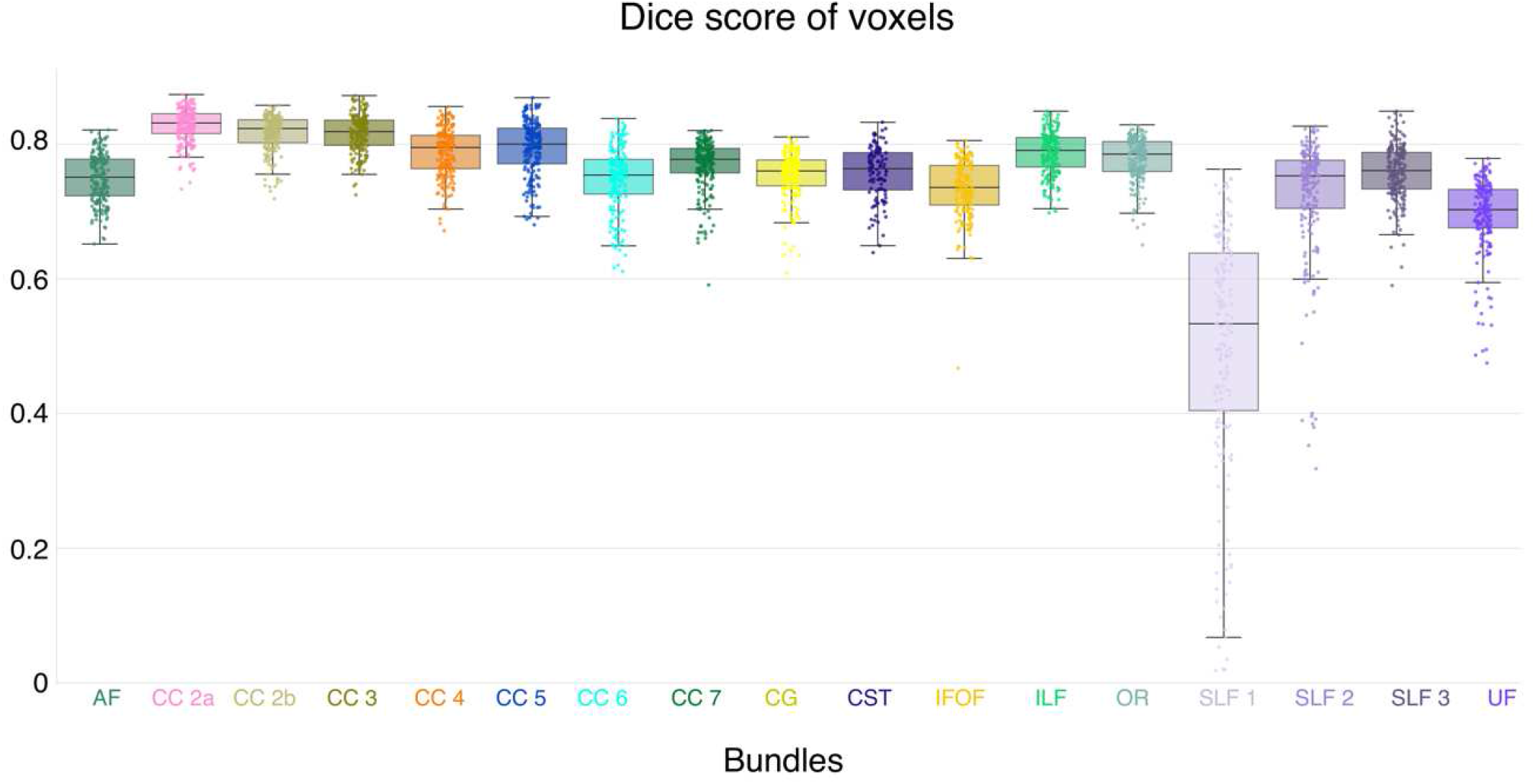
Bundles dice similarity coefficient scores for all subjects and sessions. Each dot represents one subject and session and colors correspond to each bundle.

### 3.4. Correlation analysis measures

Pearson’s correlations of all measurements across WM bundles are shown in Figure 4. This figure highlights three main aspects of the data: 1) within-model measures form a highly correlated pocket, 2) most diffusion measures are correlated, and 3) some between-model measures are correlated, while most correlations of the between-model measures are weak. More precisely, within the DTI model, the measures of MD and RD show the strongest association with each other (mean across bundles, r =0.97) and a lower association with FA (r = -0.72 and r = 0.86, respectively) and AD (r = 0.82 and r = 0.64, respectively). MD and RD also show strong associations with ECvf and ICvf (r > 0.8). On the other hand, AD (except for RD and MD), OD and the two measures of HARDI seem uncorrelated, either between or within a model (r<0.6, r<0.5, r<0.6, respectively). MTI measurements show a strong association between ihMTdR1sat and MTsat (r=0.88), ihMTR (r=0.81); and much weaker correlation between ihMTdR1sat and MTR (r=0.45). MTsat appears to be the only MTI measure that correlates, albeit moderately, with diffusion measures (r ranging from 0.3 to 0.6). This weak correlation between MTI measures and diffusion measures shows that these are sensitive to different microstructural features.

**Figure 4.**
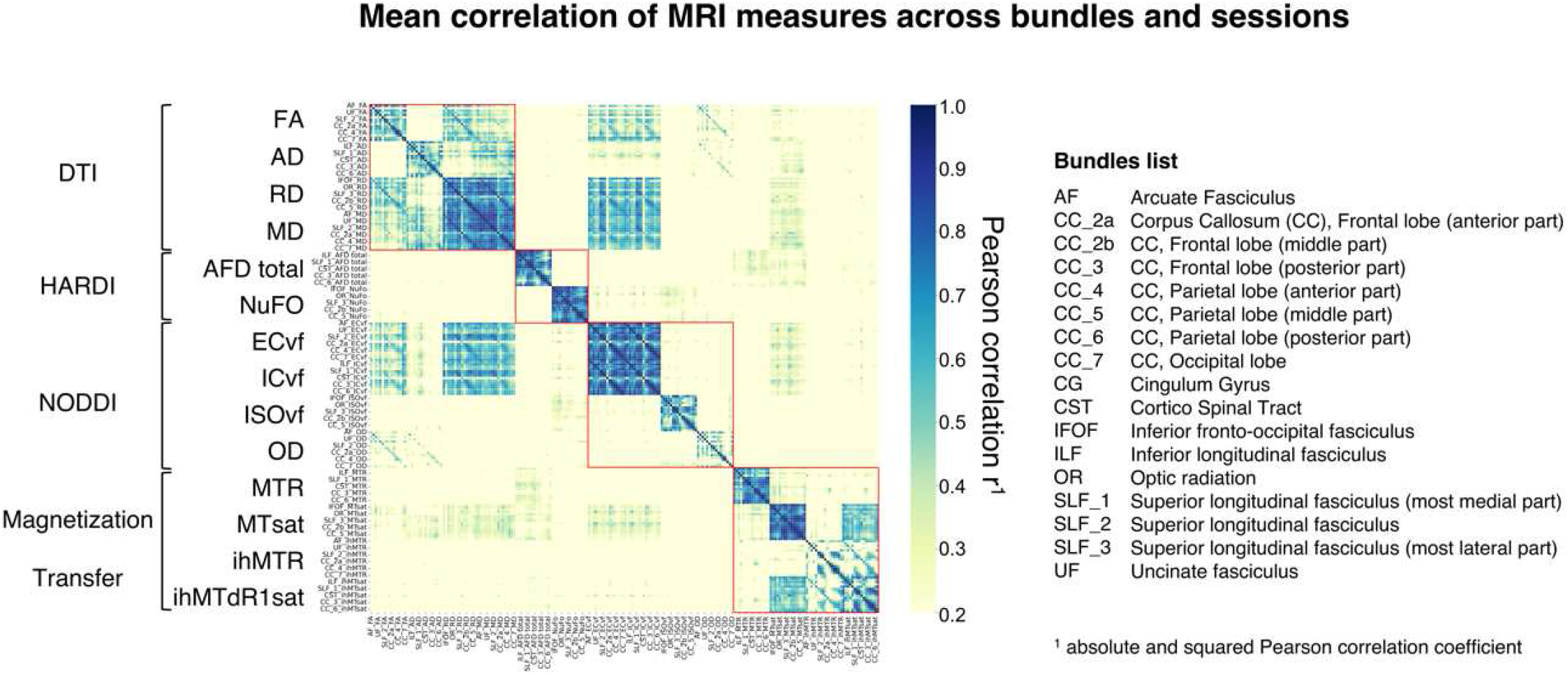
Pearson’s correlation coefficients among all MRI measures and bundles. The red squares separate each DWI model and MTI. Each measurement is ordered according to the bundle list provided in the right panel of the figure. See https://high-frequency-mri-database-supplementary.readthedocs.io/en/latest/results/correlation.html for all correlation interactive maps.

Based on these results, subsequent analyses will present the following measurements: RD, AFD total, NuFO, ISOvf, MTR and ihMTdR1sat were extracted from the 4 selected bundles: Arcuate Fasciculus (AF), section 3 of the Corpus Callosum (CC), and Cortico-Spinal Tract (CST). These measures were selected either because they show strong correlations with other measures and may share overlapping information which can cause redundancies, or because, in the opposite case, they show a pattern of decorrelation with other measures and may therefore provide different information (Cercignani & Bouyagoub, 2018; Chamberland et al., 2019). The distribution of selected measures for each subject and session is shown in Figure 5 and other measures are available at https://high-frequencymri-database-supplementary.readthedocs.io/en/latest/results/measure.html#whole-bundlemeasures.

**Figure 5.**
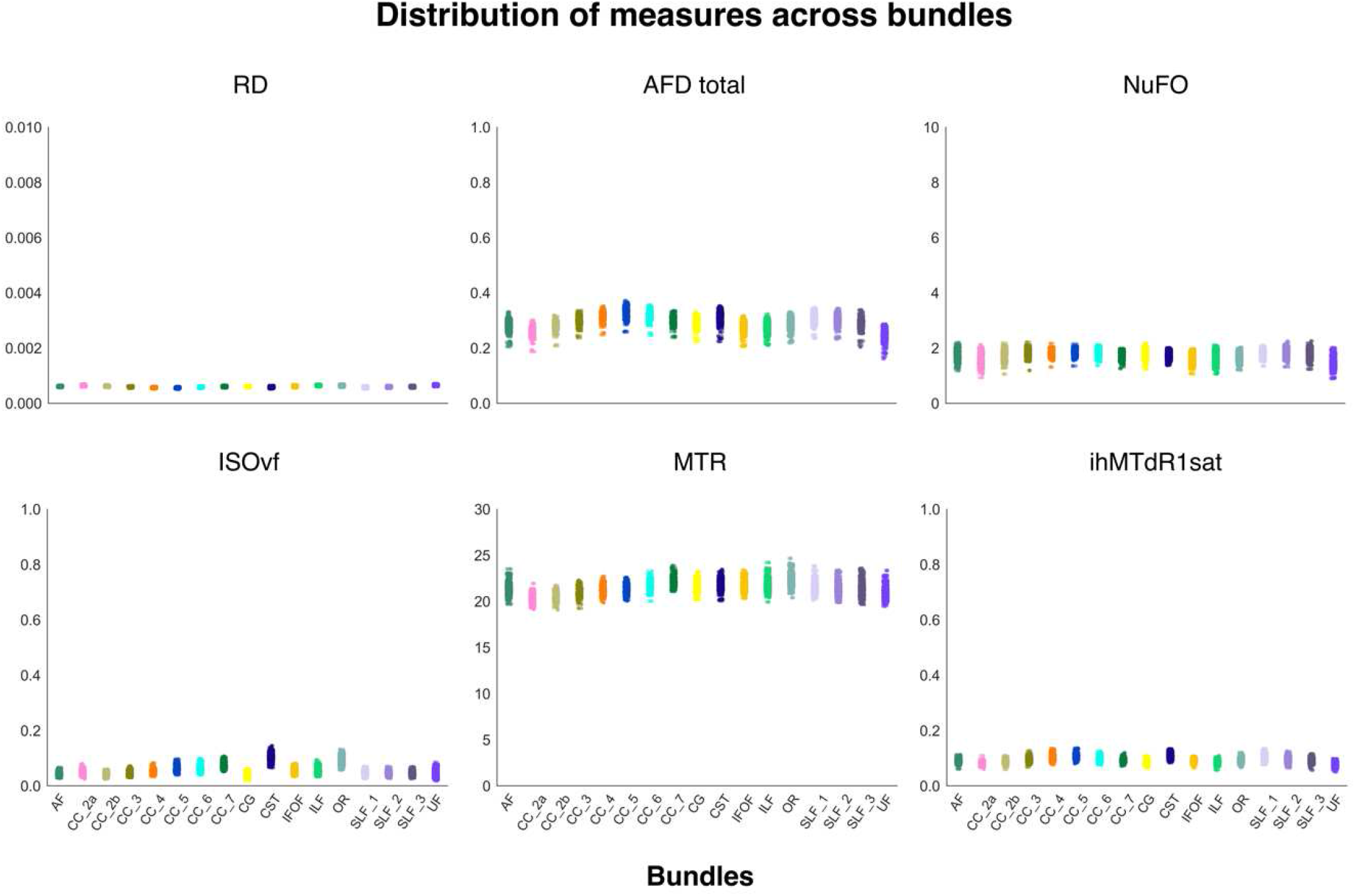
Individual measures for each bundle. Each dot represents one subject and session, and each plot represents one bundle. The colors correspond to the bundles.

### 3.5. Consistency of bundle-averaged measures

All consistency and MRI measurements are available at https://high-frequency-mri-database-supplementary.readthedocs.io/en/latest/results/consistency.html and, confidence intervals and p-values for selected bundles are shown in Supplementary Table 2. Across DWI measures, most bundles exhibit a high degree of reliability with an ICC ranging from 0.55 for NuFo to 0.93 for FA and overall low variability ranging from 1.2% for FA to 14% for ISOvf (Figure 6). As expected, DTI measures showed consistently the highest reliability (95 % of ICC were higher than 0.80) and lowest variability (90 % of CVw and CVb were lower than 5 %) with higher CVb compared to CVw and small variation across bundles (Figure 6). For HARDI measures, AFD total showed high reliability (ICC > 0.7 [0.73-0.8]) and low variability (CVw < 4.2% [1.8%-4%] and CVb < 6% [2.3%-5.7%]) for all bundles. NuFO showed lower reliability (ICC ∼ 0.62 [0.55 - 0.7]) and higher variability (CVw ∼ 8.1% [5.7%-11%] and CVb ∼ 11.5% [9%-14%]) compared to AFD total (Figure 6). NODDI measures showed lower reliability and higher variability compared to DTI measures. However, in general, OD showed consistently high reliability (ICC>∼ 0.75) and low variability (CVw < 9% and CVb < 15%) followed by ICvf and ECvf with good reliability (ICC > 0.7) and variability (CVw and CVb <10%) across all bundles. ISOvf measure represents the lowest reproducible measure of all NODDI maps with moderate reliability (ICC∼ 0.64) and greater variability (CVw and CVb <16%) (Figure 6).

**Figure 6.**
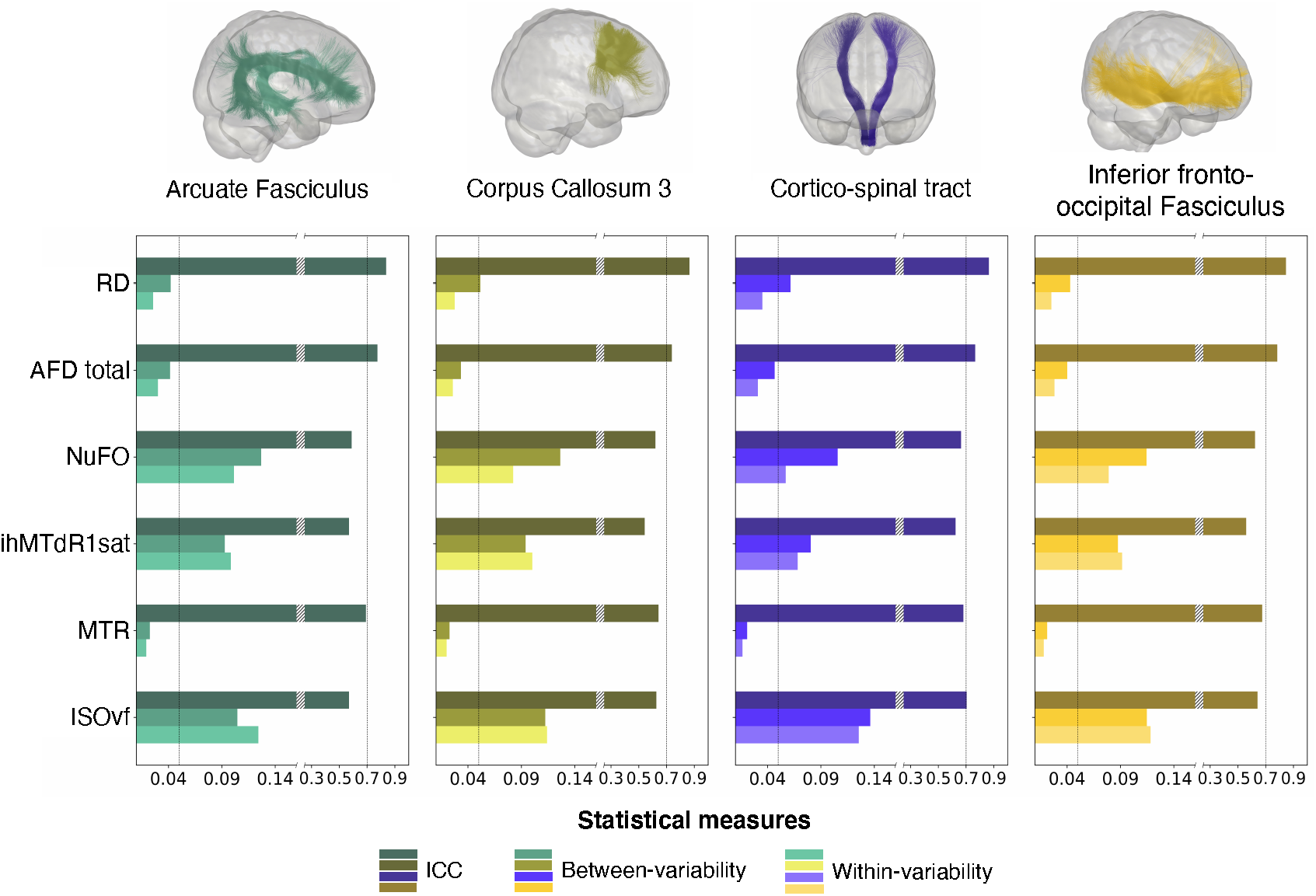
Consistency of bundle-averaged measurements of the selected bundles and measures. Each bar represents a measure of consistency with a decreasing saturation: the value of ICC in dark colors (the higher the better), Between-variability in medium colors (the lower the better) and within-variability in light colors (the lower the better). A break in the x-axis has been introduced to facilitate reading and the ICC and variability scales have been adapted.

Compared to DWI measures, a more variable pattern of results emerged for MTI measures with ICC ranging from 0.36 to 0.84 and CVs ranging from 1.4% to 13.8% across bundles. MTsat measures showed consistently the highest reliability (ICC> 0.7 except for UF and CC2a) followed by MTR (ICC∼ 0.67 across bundles). Both MT measures showed the lowest variability with CVs < 5% for all bundles. Noted that MTsat showed higher variability (CVw ∼ 2.8% and CVb ∼ 4%) compared to MTR (CVw ∼ 1.9% and CVb ∼ 2.1%) (Figure 6). In contrast, ihMT measures showed lower reliability with a mean ICC of 0.53 [0.37-0.67] and higher variability with a mean CVb of 7.9% [4.6%-11.8%] and CVw of 8.9% [6% - 13.8%] compared to MT measures. As for the MT measures, ihMTdR1sat exhibited higher ICC but higher variability compared to ihMTR (Figure 6).

### 3.6. Consistency profiles along bundles

The distribution of measures along the bundle is https://high-frequency-mri-database-supplementary.readthedocs.io/en/latest/results/measure.html#profile-bundle-measures. Globally, whatever the model and the measurement, the consistency measures quantified along the bundles display good stability or a low variability of their profile depending on the bundles (Figure 7). The different parts of the CC consistently show more variable profiles depending on the bundle sections, while the SLF bundles show the most stable profiles. On the other hand, unlike the whole bundle consistency measures, the values of ICCs profiles are lower, and the values of CVs profiles are higher (Figure 7, see also https://high-frequency-mri-database-supplementary.readthedocs.io/en/latest/results/consistency.html#profile-bundle-consistency).

**Figure 7.**
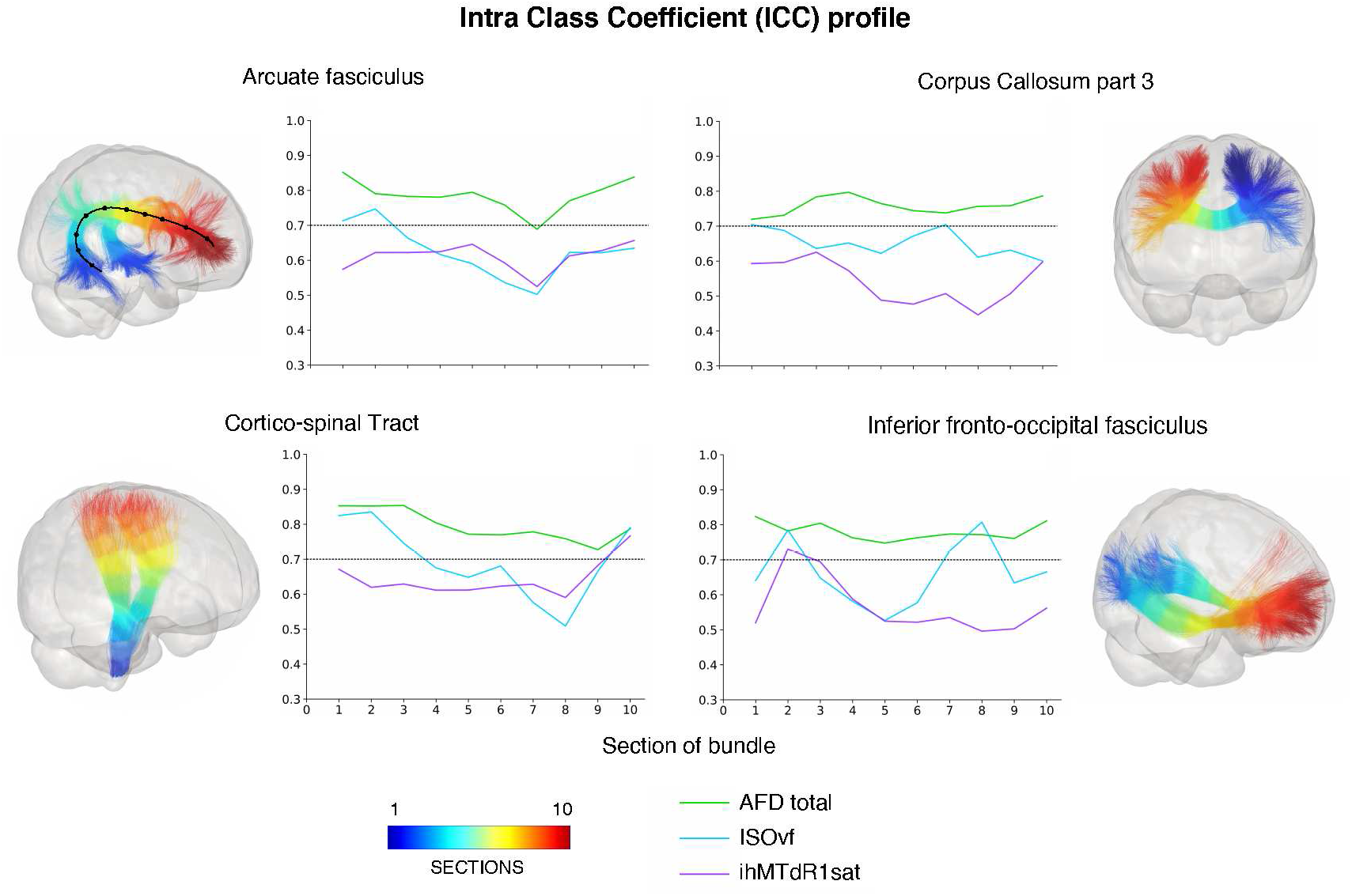
Consistency profile for selected bundles and three measures. The colors displayed on the bundles represent the section numbers from 1 (blue) to 10 (red) corresponding to the graphs. Each line of the graphs represents a measure with AFD total in green, ISOvf in light blue and ihMTdRlsat in purple. Only these three measurements for ICC results are displayed to facilitate the results’ reading and represent the different profiles observed. On the AF bundle, we show a black line with dots representing each of the 10 track-section.

More precisely, similarly to whole bundle results, consistency of DTI measures quantified along the bundles most often displays very stable profiles with higher ICCs (mean across section = 0.8 [0.64-0.95]) and lower CVs values (CVb: 4.5% [0.8%-13%], CVw: 2.8% [0.7%-9.5%]) with respect to other measures (Figure 7). Consistency profile of HARDI measures show analogous patterns to the DTI-derived ones, with similar ICCs and CVs values for AFD total (Figure 7, green line) and lower ICCs and higher CVs values for NuFO (ICC:0.72 [0.5-0.88], CVb: 6.2% [1.1%-14.1%], CVw: 4.4% [0.9%-14.5%]).For NODDI measurements, ICCs and CVs profiles show more variability along the bundles (Figure 7), but overall tend to be rather high to moderate for ICCs (0.76 [0.5-0.95]) and low for CVs, with mean value across the section of 8% [0.9%-28%] and 6.4% [0.8%-21%] for CVb and CVw respectively (Figure 7). Again, OD shows stable and higher consistency measurements in contrast to ISOvf whose consistency profile is more variable (Figure 7, light blue line).

Regarding the MTI measures, the consistency exhibits globally stable profiles for most bundles, with more stable CV profiles than ICC profiles. As for the whole bundle results, the highest ICCs and lowest CVs values are found for MTsat and MTR measures, while ihMT measures showed lower reliability (ICC ∼ 0.51 [0.3 - 0.82]) and higher variability (CVw ∼ 7.6% [3.1%-15.6%] and CVb ∼ 6.5% [1.9%-15%]) (purple line in Figure 7).

### 3.7. Impact of fiber population on consistency

For all measures, the bundle compartmentalization into “single” and “multi” fiber population regions affects ICCs and CVs values regardless of the measure (Figure 8, Supplementary Table 2, https://high-frequency-mri-database-supplementary.readthedocs.io/en/latest/results/fiber_population_consistency.html#whole-bundle-consistency). Compared to ICC computed from average bundle, the compartmentalization revealed higher ICCs (mean across bundles, ICC: 0.72, 0.82 and 0.81 for average, multi- and single-compartment respectively) and, equal to lower CVs across bundles (mean across bundles, CVb/CVw: 6.7%/5.2%, 3.7%/2.7% and 4.6%/3.4% for average, multi and single-compartment respectively). More precisely, ICCs measures from multi-compartment were higher, while the ICCs value observed for single compartment is lower and inversely for CVs (see https://high-frequency-mri-database-supplementary.readthedocs.io/en/latest/results/fiber_population_consistency.html#whole-bundle-consistency). Differences between single and multi compartments can be moderate, especially for DTI measurements, or more important such as MTI measurements. Some measures exhibit a different pattern than others. This is the case for the OD measurement of the NODDI model, whose reliability pattern is inverted. As expected, since OD is a measure of the dispersion of fiber orientation in the voxel, a higher or equal ICCs to the average for single compartment compared to multiple compartments for most bundles is not surprising. This inverted pattern is also found for ihMTdR1sat and MTsat measures for the CST bundle (Figure 8, purple line). In addition, the impact of compartmentalization has a more moderate effect on the consistency of SLF bundles measures. Indeed, the values of ICCs and CVs are generally equal (or present a slight difference) between the two compartments. This moderate effect is also found for DTI measures, for most, but not all, bundles (Figure 8, see RD as an example, red line).

**Figure 8.**
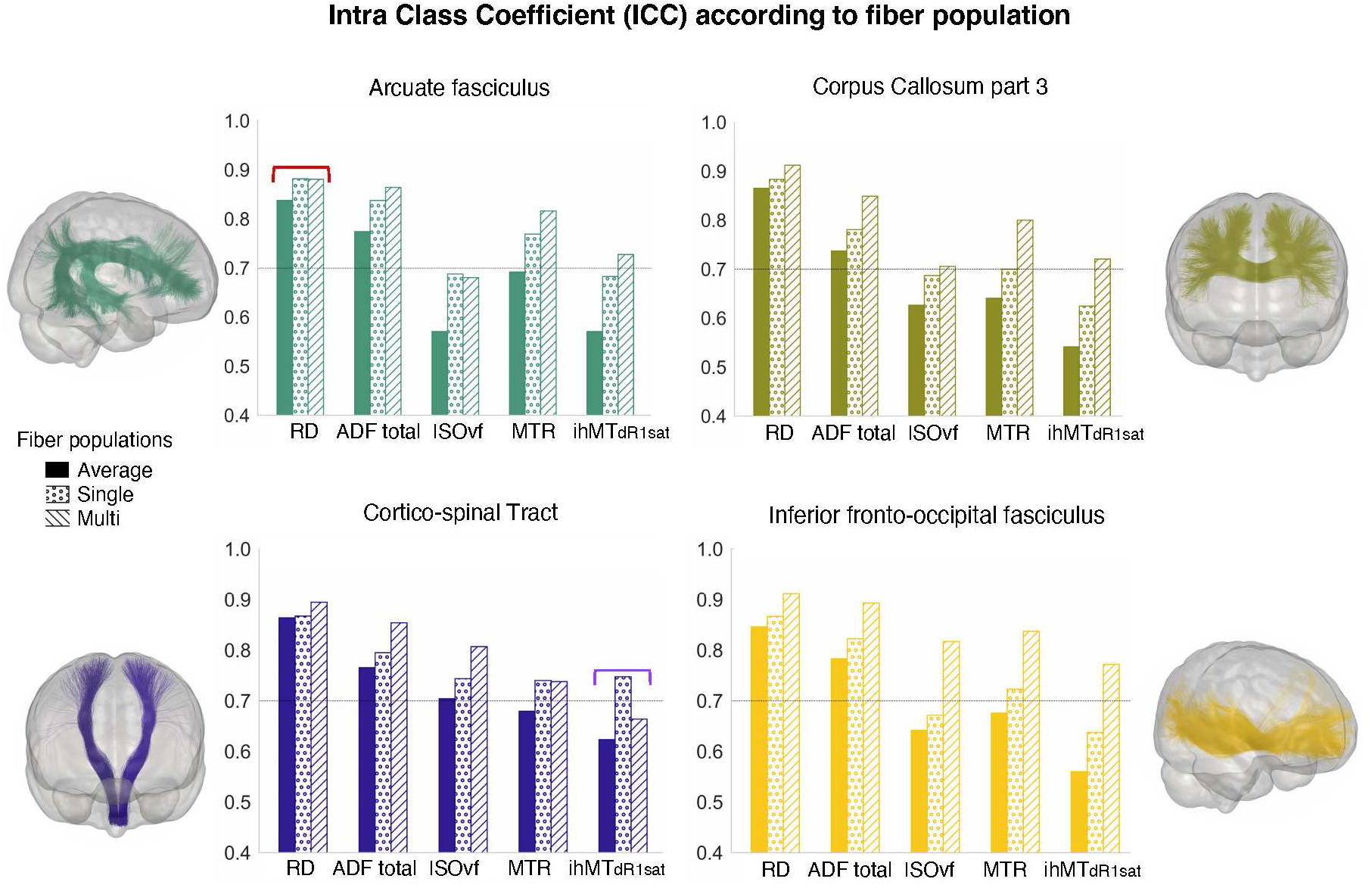
Impact of fiber populations on bundle-averaged consistency values for selected bundles. The colors correspond to the bundles. The bars represent the ICCs values, full for “average”, dotted hatch for “single” and diagonal hatch for “multi” compartment.

The compartmentalization also affects the mean values of the measurements. The mean of the measures in the multi-compartment was lower than those found in the single-compartment (for example, mean CST FA: 0.47, 0.53 and 0.35 for average, single- and multi-compartment respectively). Globally, there is a decrease (or increase depending on the measure) in the mean values of the measures with increasing fiber populations. The distribution profile of measures is available https://high-frequency-mri-database-supplementary.readthedocs.io/en/latest/results/fiber_population_measures.html#profile-bunlde-measures. Figure 9 presents profile examples for some measures and the associated consistency values for CST bundle. The compartmentalization of the bundle profiles into “single” and “multi” fiber population regions showed a similar effect to the results on the whole bundle profiles, with higher multi-compartment ICCs compared to the single-compartment and, inversely for CVs (Figure 9, https://high-frequency-mri-database-supplementary.readthedocs.io/en/latest/results/fiber==12opulationconsistency.html#profile-bundle-consistency). The single-compartment profile is generally close to or equal to the whole bundle profile, while the multi-compartment is more distant from the average profile. Some bundles present a different pattern for the inferior (section 1-3) and superior (section 8-10; or anterior/posterior) sections compared to middle sections. For example, CST exhibits a higher ICC in the single compartment for inferior sections (1-2), whereas the superior sections (8-10) show weak differences between the compartments compared to the middle part of the bundle (sections 3-7, Figure 9). Again, SLF bundles show globally the most stable profiles with a low impact of compartmentalization on consistency, whereas the CC bundles show the reverse pattern. Finally, this compartmentalization effect is also found in the measurement profiles (Figure 9). This effect is greater for some measures such as FA or OD while others are less affected such as ISOvf, ICvf, MD or MTR. Like consistency results, the compartmentalization does not uniformly affect the measurement profiles of bundles according to the sections, with a different effect between the sections in the middle and those at the ends of the bundle (Figure 9).

**Figure 9.**
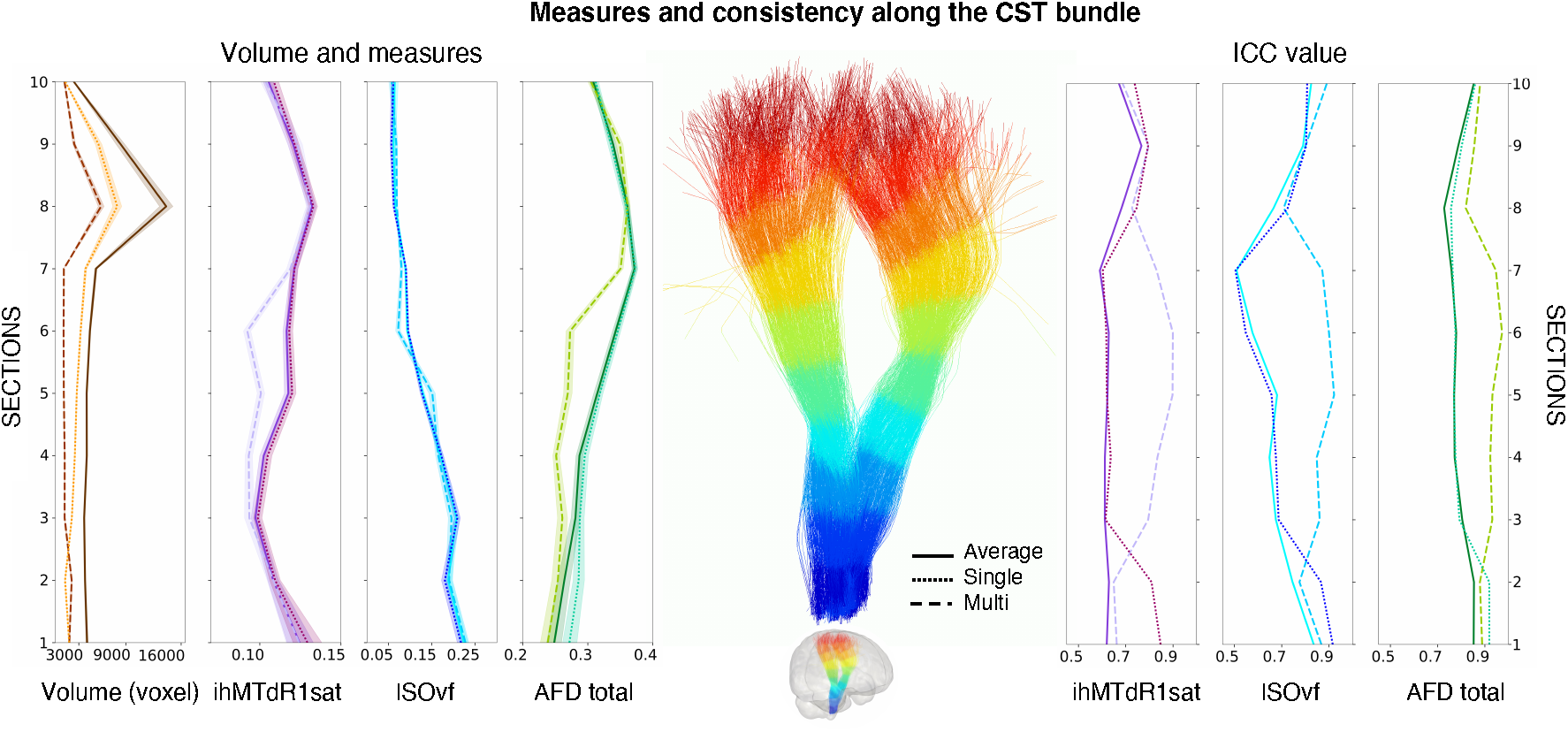
Volume, measures and consistency values per section and compartment of the CST bundle. The left panel shows volume and measures, the right panel shows ICC value. Continuous line represents values obtained from bundle average, dashed line represents value from multi compartment and dot line represents value from single compartment. Colors correspond to measures, volume in brown, ihMTdRlsat in purple, isoVF in light blue and AFD total in green.

## 4. Discussion

Using an optimized “high-frequency” repeated-measure study collected from twenty healthy subjects, we assessed the consistency of multiple WM microstructural measures across the bundles, with special emphasis on the most frequently used image analysis approaches. The Dice scores reveal a good spatial agreement of the segmentation of the bundles across the sessions, except for SLF 1 whose results should be taken with caution. The results show that the reliability and variability of DWI measures are good (ICC > 0.7; CVw and CVb < 15%) across the bundles and especially excellent for most DTI measures as well as AFD total and OD index from the HARDI and NODDI models, respectively (CVw and CVb < 5%, ICC > 0.75). In addition, the profile consistencies of these measures are comparable to the whole bundles, with voxel values well above ICC > 0.7 and CVw < 4% along the bundle. In contrast, MTI showed good reliability and variability for MT measurements (CV < 8%, ICC ≥ 0.7) and moderate for ihMT measurements (CV < 15%, ICC ≥ 0.5). We also showed that the number of fiber populations affects the consistency of the measurements, with a moderate effect on the DTI and HARDI measurements, and a more important effect on the NODDI and MTI measurements. Finally, SLF 2-3 and CC 4 to CC 6 bundles showed the most consistent MRI measurements followed by AF, CST, CC 7, OR, CC3, IFOF and CG, while CC 2a, CC 2b, UF and ILF have more moderate consistency measurements (Figure 10).

**Figure 10.**
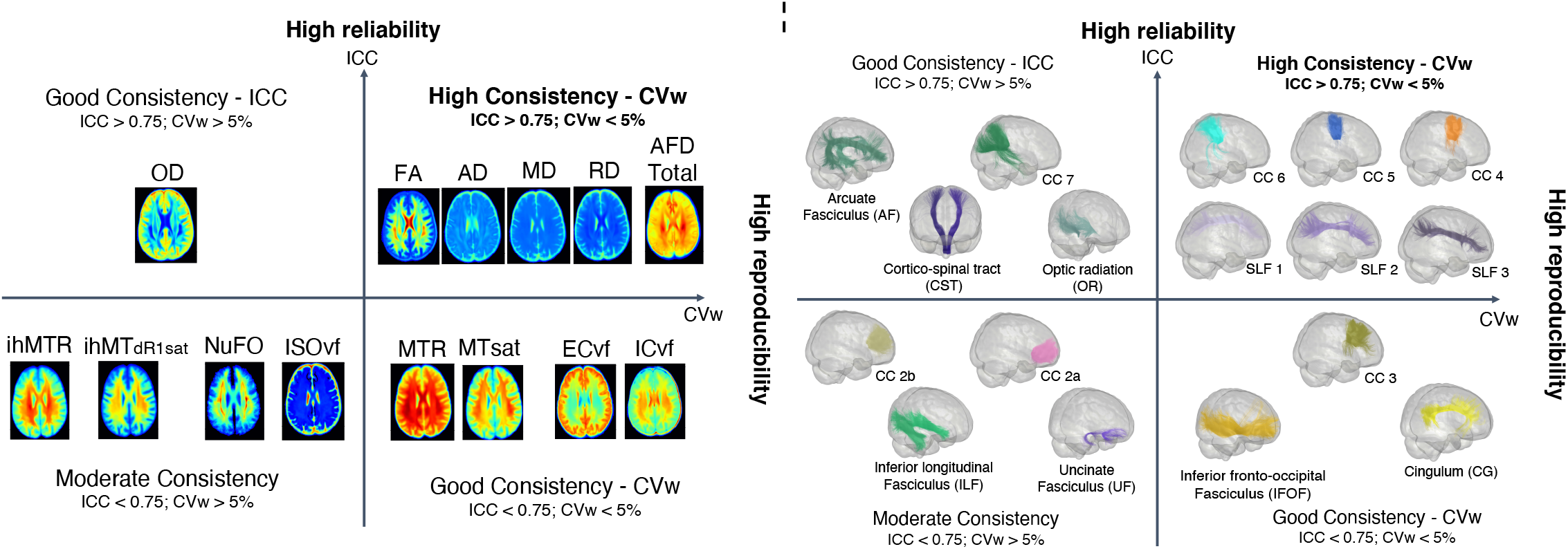
Classification of bundles and measures into four groups according to ICC values and within-variability (CVw). The x-axis represents CVw values (i.e., reproducibility), and the y-axis represents ICC values (i.e., reliability). High consistency group represents low CVw value < 0.05 (5%) and high ICC value > 0.75 (high reliability); Good consistency - ICC: high ICC value > 0.75 but high CVw value > 0.05 (>5%); Good consistency - CVw: low CVw value < 0.05 but low ICC value < 0.75 and finally, Moderate consistency: high CVw value > 0.05 (>5%) and low ICC value < 0.75.

We chose a sample of techniques including popular and novel MRI techniques to do all the consistency analyses. However, many other MRI techniques and processing tools are available to do similar tasks and many ways to assess reliability and variability. For instance, there are different methods for performing tractography (Maier-Hein et al., 2017; Yendiki et al., 2011) or identifying bundles (Schilling et al., 2021). These latter parameters could influence both variability and reliability. Additionally, several other microstructural measures can be characterized. The intent here was not to provide an analysis between different processing tools or parameters, and therefore we do not recommend using the consistency values from this study as an absolute value. Instead, we aimed to contribute to a global understanding of the variability and reliability of popular WM imaging techniques such as DWI as well as newer techniques such as ihMT.

### Diffusion consistency measures of whole and along-bundle profiling

Here, we demonstrated good to excellent consistency, for both the whole bundle and along the bundle, of most diffusion measures used in MRI studies. More precisely, the reliability and variability of DTI measurements are comparable or superior to the previous DTI studies, with ICC above 0.7 and within- and between-CV around 5% and 10% (Acheson et al., 2017; Grech-Sollars et al., 2015; Hakulinen et al., 2021; Luque Laguna et al., 2020; Magnotta et al., 2012; Palacios et al., 2017; Shahim et al., 2017; Veenith et al., 2013; Zhou et al., 2018). In line with past studies, across white matter bundles, within- and between-subject CV of FA (∼5% [3%-6.5%] and ∼9.8% [6.5%-12.5%], respectively) was larger than other DTI measures, such as MD (∼1.8% [0.2%-2.4%] and ∼2.6% [1.8%-3.5%], respectively) (Grech-Sollars et al., 2015; Luque Laguna et al., 2020; Veenith et al., 2013). Moreover, previous studies report similar or lower variability for FA with CV ranging from 1% to 6% (Acheson et al., 2017; Grech-Sollars et al., 2015; Luque Laguna et al., 2020; Palacios et al., 2017; Veenith et al., 2013), while reports of MD are more variable, ranging from 2 to 7 % with most studies being around 2% within- and/or between-subject CV (Grech-Sollars et al., 2015; Magnotta et al., 2012; Shahim et al., 2017; Zhou et al., 2018). The AFD total measure derived from HARDI exhibits similar consistency to DTI with CV < 10% and ICC > 0.7. Regarding NuFO, only one study has assessed the reliability of this measure in a test-retest study and reports moderate reliability with an ICC of approximately 0.6 across four bundles (Boukadi et al., 2019). In agreement with this study, we report an ICC value of ∼0.62 ranging from 0.5 to 0.7 for all bundles. We also report high variability with a mean CVb of 11% and CVw of 8.1% ranging from 5.7% to 14%. Together, this suggests that there is a need for further validation of this measure before adopting it in longitudinal studies. On the other hand, even though NODDI measurements are inherently noisier than DTI measurements for white matter modelling - likely due to a more complex model and requiring high b-value data - within- and between-variability for ICvf and ECvf were lower or comparable to the FA variability in most bundles (Andica et al., 2020; Chung et al., 2016; Lucignani et al., 2021). However, this sensitivity to noise inherent to NODDI could explain the higher within-variability of ISOvf and OD. Indeed, despite the additional thresholding applied on ISOvf, this measure systematically presents the largest CV in all the bundles, which is consistent with studies that suggest that ISOvf is a poorly reliable parameter (Tariq, 2013, 2018). Finally, our findings of overall greater CVb for most measures compared with their corresponding CVw are in accordance with other DTI- and NODDI-based reliability studies (Andica et al., 2020; Chung et al., 2016; Lucignani et al., 2021; Tariq, 2013; Veenith et al., 2013). This result is expected, as there will be greater microstructural heterogeneity in a localized region across a population compared with multiple observations within the same subject.

### MTI consistency measures of whole and along-bundle profiling

Previous studies that include MTR reported ICCs ranging from 0.5 to 0.9 and variability <6% in the different brain regions (Schwartz et al., 2019; van der Weijden et al., 2021) which is consistent with our MTR findings although our MT measurements were derived from ihMT images. We also showed that MTsat had the highest consistency among MTI measurements with variability <5% and ICC of 0.77. These results are consistent with the only study that reported a measure of between-variability ranging from 6 to 8% (Weiskopf et al., 2013). For ihMT, we reported a mean ICC of 0.5 ranging from 0.4 to 0.56, a lower value than the previous multicenter MRI study that reported an ICC above 0.7 (L. Zhang et al., 2019, p. 20). Mchinda and colleagues’ study reports standard deviations of between-individual ihMTR under 10% (Mchinda et al., 2018; L. Zhang et al., 2019), whereas we reported lower between-subject CV (CVb < 6.8%). For ihMTdR1sat, we report moderate ICC of ∼ 0.6 ranging from 0.43 to 0.48 with high CVb ∼8.7% and CVw ∼9.1%. These differences could be explained by, first the different ways to assess variability and reliability measures, then the difference for ihMT sequences with a long cosine-modulated RF pulse, without any T1D filtering in Zhang’s report instead of a train of short RF pulses with moderate T1D filtering in this study (L. Zhang et al., 2019); and Mchinda’s report an ihMTR between-subject variability based on 1.5T MRI instead of 3T MRI in this study (Mchinda et al., 2018). In addition, Varma et al. reported that ihMTR could be varied depending on different sets of saturation parameters, especially when the saturation time is in the range of 0-200 ms (Ercan et al., 2018; Varma et al., 2015). This discrepancy between studies suggests a strong effect of experimental design and local factors on the measurements and, therefore, makes it difficult to compare our results with each other. Nevertheless, we can agree that among the MTI-derived measures, the ihMT measures appear to be more variable and less reliable than the other measures. Noted that considering these measurements are derived from the same acquisition, the ihMT effect (i.e., signal-to-noise ratio, SNR) being much weaker than a standard MT effect, it is expected that the variability of ihMT will always be greater than the MT variability. However, ihMT is more specific to myelin (Duhamel et al., 2019; Prevost et al., 2018), which is a significant advantage in clinical studies. Thus, as with all new contrast, further optimization might improve the accuracy of ihMT measures and, by extension, their consistency under different experimental conditions.

### Consistency and fiber population

Results emerging from the single *versus* multi-fiber population impact show that the fiber populations affect the consistency of measures. The higher the fiber populations of a bundle (*i*.*e*., multi fibers), the higher the ICC and lower the within- and between-subject CV. Inversely, the lower the fiber populations of a bundle (*i*.*e*., single fiber), the lower the ICC and higher the within- and between-subject CV. Despite it seems like counter-intuitive results, a smaller range of values for the compartment with multiple fiber populations for most measures compared to the single compartment explains the greater consistency (CST FA range in multi: 0.34-0.42 *vs* single 0.48-0.61). Thus, the intrinsic variability of the measurements in the multi compartment is already more restricted than in the single compartment. Although the method and the variability values are not directly comparable, Volz et al. also report greater variability of the compartment with a single fiber population compared to compartments with multiple fiber populations (Volz et al., 2018). In agreement with this study, diffusion and myelin measurements *per se* are also impacted by compartmentalization, with higher value in the single compartment compared to the multiple compartments (and vice versa depending on the measurements). This reinforces the idea that the organization of the underlying WM affects both measures and consistency. Moreover, some bundles are more affected than others, this is notably the case of the CC which, unlike the SLF bundles, displays a more pronounced compartmentalization effect. Beyond the bundle, we also showed that some measurements are more affected than others, suggesting that the affected measurements are sensitive to the orientation of the fibers of the white matter. This has been described for the ihMT which shows a dependence on the orientation of the fibers with respect to B0. Thus, in the presence of several fiber populations, this effect could be averaged, thereby improving the consistency of the measurements. Together, these results suggest that the dissociation of voxels according to the number of fibers populations may be relevant and must be tested in future studies.

### What is the contribution of this study to clinical and research studies?

Assessing the consistency of measurements extracted from MRI images represents an important step toward validating this approach in longitudinal studies and clinical trials. Clinical applications favor measures with high reliability, which optimizes a trade-off between the two variability components with low within-variability (*i*.*e*., more stable across different measuring times) and high between-variability (*i*.*e*., more differentiable across participants). This variability pattern is a necessary criterion for the high validity of a biomarker, which can be used to diagnose, monitor, and predict neurological consequences by clarifying the effects of a disease or treatment.

We have shown that the measures corresponding to this criterion are the “simplest” *i*.*e*., the DTI-derived measures, particularly FA, RD, AD and MD. However, this study sheds light on more advanced measures that could be promising candidates as biomarkers. This is particularly the case of the AFD total - an axonal density index - which can be interesting in pathologies where the axons are altered and disrupted, such as Alzheimer’s disease (Roy et al., 2021). Next, a growing body of evidence shows that NODDI-derived ICvf and ECvf measures - which provide a proxy of the axonal density and the volume of extracellular space - are predictive of response to treatment (Dowell et al., 2019; Kraguljac et al., 2019; Sarrazin et al., 2019), reinforcing that these measures may be clinically relevant biomarkers. The OD measure is a dispersion measure of fiber orientation in the voxel whose reliability is good and which covaries with the NuFO (Chamberland et al., 2019). The latter displays more moderate reliability, but unlike OD, which requires a multi-shell DWI acquisition, NuFO can be extracted from clinical acquisitions without requiring advanced modelling, which is an important advantage for clinical studies. In addition, recent studies show that it is important from the biological mechanisms point of view to consider fiber orientation dispersion, especially in the crossing fibers regions such as the semioval centrum (Andersen et al., 2020; Chad et al., 2021; Mito et al., 2018; Schilling et al., 2020). A DTI study showed in MS patients that considering the orientation of the fibers for the FA highlights changes in WM related to disability, while the standard FA failed to do so (Andersen et al., 2020). Another study shows that a fixel-based analysis (D. A. Raffelt et al., 2017) reveals a specific degeneration of the SLF with preservation of the CST and the CC in Alzheimer’s disease whereas this degeneration results in an increase in FA with conventional DTI measures, *i*.*e*., the decrease in FA of the SLF bundle leads to a “virtual” increase in FA in the CST (Mito et al., 2018). This indicates that consideration of different fiber populations can detect a change in bundle-specific MRI measurements, without causing significant abnormality in other bundles that intersect in the same region (Doan et al., 2017; Lee et al., 2015; Mito et al., 2018). Therefore, appropriate consideration of different fiber orientations using specific methods or indices such as NuFO or OD could play a critical role in understanding brain disease processes where conventional measurements are “blind”.

Regarding the MTI, although the MTR is the most used measure, this study shows that the MTsat presents a lower within-variability and a greater between-variability compared to the MTR (https://high-frequency-mri-database-supplementary.readthedocs.io/en/latest/results/consistency.html#within-variability). Again, this indicates that MTsat would be a more favorable biomarker than MTR. Moreover, recent studies have shown that MTsat is more sensitive than MTR in MS (Granziera et al., 2021; Lema et al., 2017; Saccenti et al., 2020). Despite lower consistency, recent studies support the use of ihMT in clinical studies due to its specificity to myelin, especially in patients with MS (Obberghen et al., 2018; Prevost et al., 2018; Varma et al., 2015). Indeed, albeit preliminary due to the small number of subjects, a recent study showed that in MS patients ihMTR was correlated with clinical disability, whereas MTR failed to do so (Obberghen et al., 2018). Moreover, many efforts have been made recently to overcome significant technical limitations - especially on its feasibility on different scanners - and make ihMT an applicable tool in daily clinical practice that may outperform MT measures soon (O. M. Girard et al., 2015; Soustelle et al., 2022; Varma et al., 2018; Wood et al., 2020).

Regarding the bundles, our results suggest that all bundles that show high consistency, whether extracted on whole or along the bundles, are involved in different pathologies, aging or development. This is particularly the case for AF, ILF, IFOF, CST, UF and CG which have very good consistency *per se* (Dice score) (Atkinson-Clement et al., 2017; Beaudoin et al., 2021; Bergamino et al., 2020; Coelho et al., 2021). A more moderate observation can be made regarding some parts of the corpus callosum, which showed more variable consistency of data. Beyond this, the profiles reveal good to excellent levels of coherence, like the measurements from whole bundle. Interestingly, the consistency levels and measurements vary according to the sections along the bundle. This suggests that along-bundle profiling could help to reliably highlight more subtle changes such as the presence of a white matter lesion, which may impact one or more parts of the bundle without affecting the whole bundle. This is supported by a recent study which shows that AFD profiles are affected by the presence of a lesion and are significantly different at the location of WMH in fronto-pontine tracts bundle in mild-cognitive impairment subjects with high WM lesion load (T. Kim et al., 2022).

## 5. Conclusion

Using an optimized “high-frequency” repeated-measure study collected from twenty healthy subjects, we showed that the reliability and variability of DWI measures are excellent to good across the bundles as well as along the bundle. In contrast, MTI showed good reliability and variability for MT measurements and moderate for ihMT measurements. We also showed that the number of fiber populations affects the consistency of the measurements, with a moderate effect on the DTI and HARDI measurements, and a more important effect on the NODDI and MTI measurements. Finally, the most consistent MRI measurements are found for SLF 2-3 and CC 4 to CC 6 bundles, then for AF, CST, CC 7, OR, CC3, IFOF and CG, while CC 2a, CC 2b, UF and ILF have more moderate consistency measurements.

## Acknowledgments

We would like to thank the participants who accepted to take part in this study. We also thank the radiology team at Centre Hospitalier Universitaire of Sherbrooke (CHUS) for their support with the scanning of participants.

## 6. Data availability

The raw and generated datasets during the current study are not publicly available because we are not the exclusive owners of the data. However, they are available from the corresponding author under reasonable request and data sharing agreements with the owners.

## 7. Code availability

All Nextflow and code used to process the T1 and DWI images, ihMT images, RecoBundlesX and Tractometry are available at https://github.com/scilus. Operations like generating and eroding masks were done using Scilpy scripts available at https://github.com/scilus/scilpy. Consistency analysis code is available at https://github.com/AlexVCaron/longitudinal_image_statistics.

## 8. Author contributions

ME drafted the manuscript, contributed to the design of the study, reviewed the literature, and contributed to data collection, analyzed, and interpreted the data. M. Descoteaux designed the study, supervised data analysis and interpretation, and contributed to drafting the manuscript. M. Descoteaux and GG contributed to MRI data acquisition. AT contributed to the design of the study and data collection. ME, GT, and GG contributed to developing the inhomogeneous magnetization transfer pipeline. AVC, GT and M. Dumont helped with the data analysis and AVC developed the voxel-level consistency analysis script. JCH, GT, LM and M. Dumont contributed to image processing. FR performed the bundle’s consistency analysis. All authors revised the final version of the manuscript. SM designed the study, data interpretation, and review the manuscript. MB and FS helped with the data interpretation and review the manuscript.

