## Supplementary data for "High-frequency longitudinal white matter diffusion- & myelin-based MRI database: reliability and variability"

### 1.1 Bundle sections

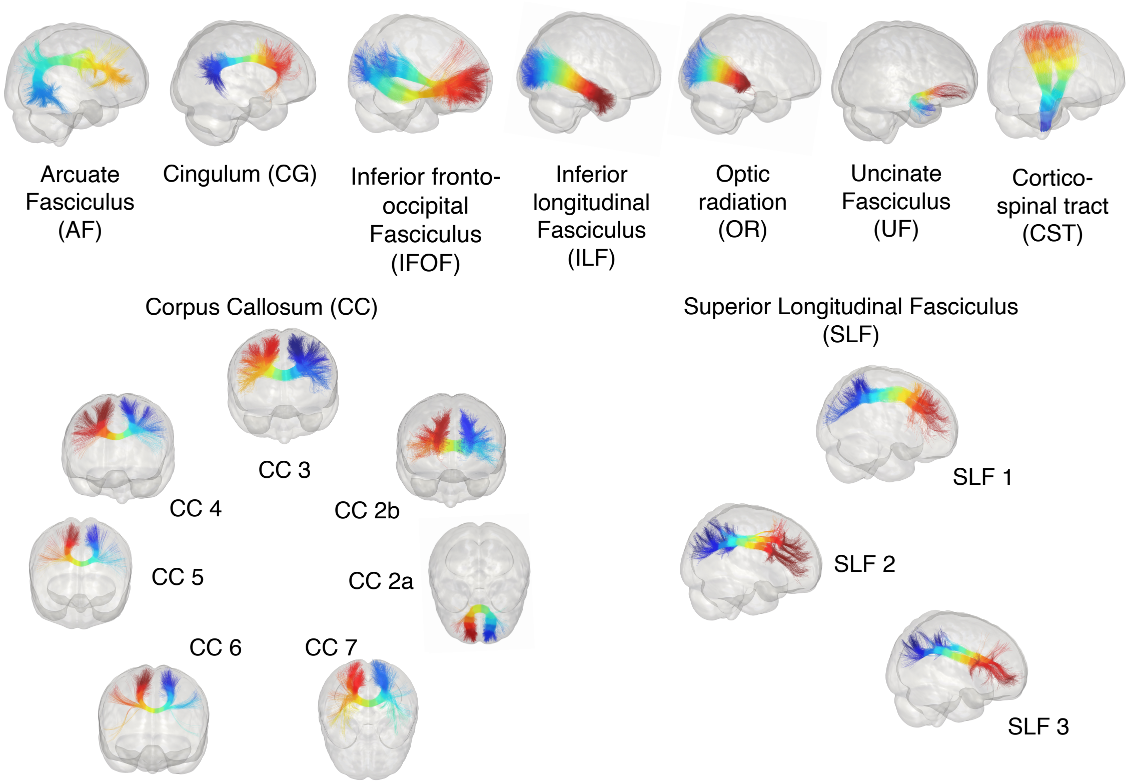


**Figure 1 Representation of bundle sections.** The major bundle models used by RecobundlesX as shape priors to extract the bundles from the whole tractogram were resampling into 10 segments for illustration. Left and right have been merged. The colors displayed on the bundles represent the section numbers from 1 (blue) to 10 (red).

### 1.2 Pipeline analysis overview description

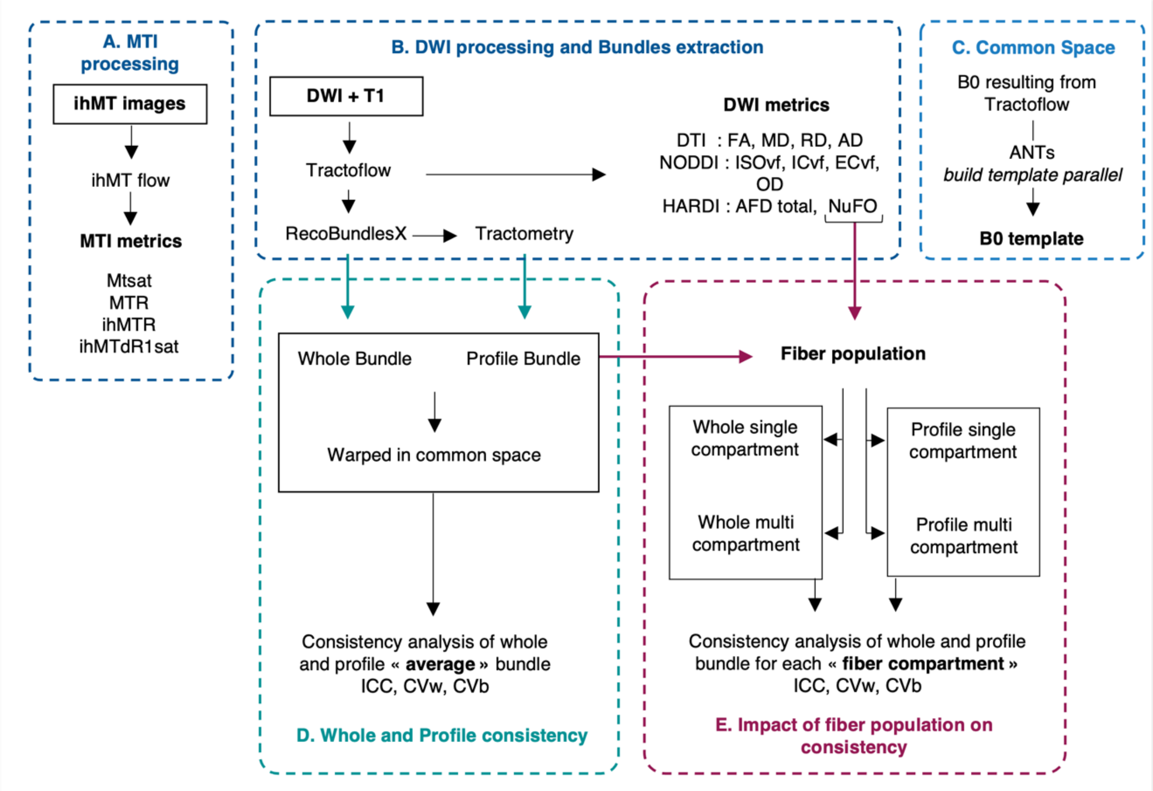


**Figure 2 Processes and analyses overview.** In A (blue), MTI input and output files using ihMT flow. In B (blue), the DWI process using Tractoflow to generate the tractograms and diffusion measures maps, the bundles' virtual segmentation using RecoBundlesX to obtain whole bundle mask, and bundles resampling using the Tractometry flow which provides bundle profile masks. In C (light blue), diffusion common space generation using ANTs. In D (green), consistency analyses of diffusion and myelin measures from the whole and profile bundle masks. In E (red) processes that take the NuFO map generated in B and the whole and profile bundle mask generated in D to separate them into single and multi compartments masks and perform consistency analyses of diffusion and myelin measures from these masks.

### 1.3 Impact of resampling on bundles volume

To ensure that each section of bundles contains enough voxels to assess consistency measurements, we extracted the volume of each section corresponding to the bundle profile analyses. A minimum threshold of 1000 voxels (dotted line) was used to perform analyses, therefore, sections of bundles with fewer than 1000 voxels were excluded. Only the volume of section 10 of the cingulum is below the threshold (circle and red arrow). This section was therefore excluded from the analyses.

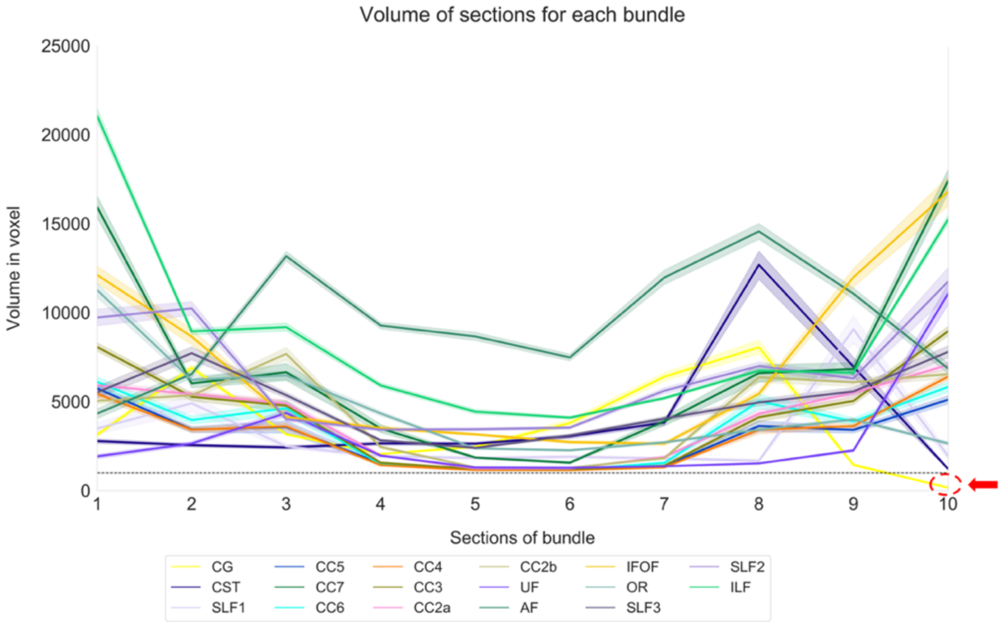


**Figure 3 Volume corresponding to each section for each bundle.** Colors code for bundles. The volume is expressed in voxels. The black dotted line corresponds to the threshold of 1000 voxels.

### 1.4 Impact of ISOvf thresholding on consistency measures

Isotropic Volume Fraction (ISOvf) has generally very low values in the WM and many studies emphasized the poor reliability of this NODDI parameter. To improve the consistency of this parameter, we evaluated the impact of different thresholds to remove values close to zero. A range of thresholds between 0 and 1 with a step size of 0.1 was used. The threshold of 0.045 was chosen because it corresponds relatively to the inflection point of the ICC curve (red dotted line).

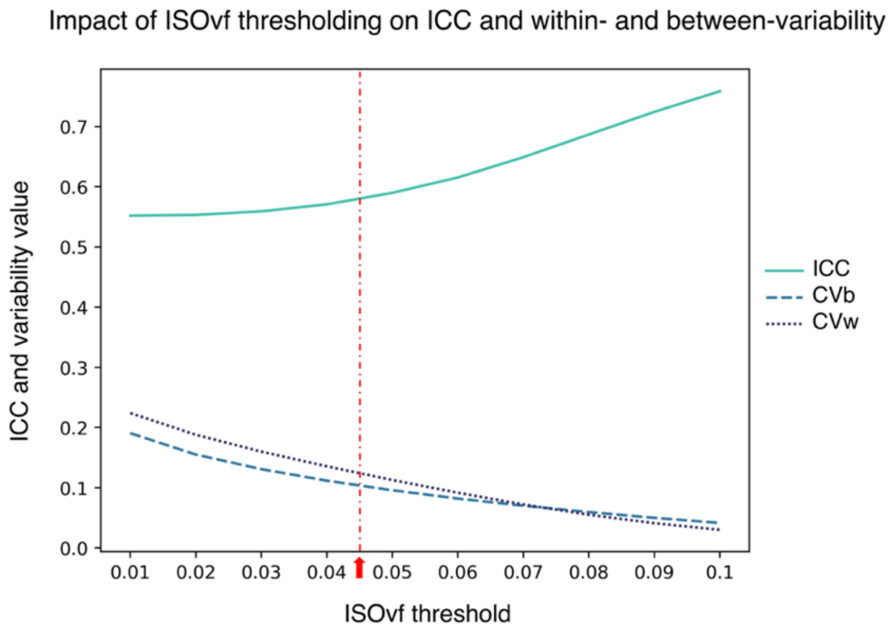


**Figure 4 Consistency measures according to ISOvf thresholds.** The graph illustrates the ICC (green), within- (dark blue, dotted line) and between-variability (blue, dashed line) values according to the different ISOvf thresholds. The red dotted line represents the chosen threshold of 0.045.

**Supplementary Table 1 Descriptive and consistency metrics for MRI measurements.**

For AD, RD, and MD units of Mean, SD and Range are $10^{-3}$ $mm^2/s$; CVb = averaged between-subject, CVw = averaged within-subject, ICC = averaged across sessions, *** $p < 0.001$, ** $p<0.01$, *$p<0.05$.

|  | Model | Measures | Mean | SD | Range (min-max) | CVb | CVw | ICC | ICC CI |
|---|---|---|---|---|---|---|---|---|---|
| AF | DTI | FA | 0.368 | 0.015 | 0.055 | 0.008 | 0.033 | 0.917*** | 0.911-0.921 |
|  |  | AD (10-3) | 1.032 | 0.019 | 0.085 | 0.033 | 0.016 | 0.902*** | 0.893-0.907 |
|  |  | RD (10-3) | 0.585 | 0.020 | 0.074 | 0.037 | 0.018 | 0.903*** | 0.890-0.914 |
|  |  | MD (10-3) | 0.734 | 0.018 | 0.067 | 0.019 | 0.011 | 0.853*** | 0.822-0.872 |
|  | HARDI | AFD total | 0.283 | 0.018 | 0.105 | 0.031 | 0.019 | 0.820*** | 0.794-0.866 |
|  |  | NuFO | 1.228 | 0.043 | 0.256 | 0.102 | 0.067 | 0.685* | 0.674-0.702 |
|  | NODDI | ECvf | 0.438 | 0.024 | 0.093 | 0.044 | 0.035 | 0.793** | 0.766-0.817 |
|  |  | ICvf | 0.562 | 0.024 | 0.093 | 0.029 | 0.020 | 0.792** | 0.768-0.821 |
|  |  | ISOvf | 0.051 | 0.008 | 0.040 | 0.104 | 0.124 | 0.571* | 0.548-0.587 |
|  |  | OD | 0.294 | 0.011 | 0.052 | 0.108 | 0.050 | 0.913*** | 0.907-0.918 |
|  | MTI | MTR | 22.000 | 0.569 | 3.187 | 0.017 | 0.014 | 0.738** | 0.643-0.783 |
|  |  | MTsat | 3.520 | 0.147 | 0.974 | 0.054 | 0.061 | 0.569* | 0.515-0.604 |
|  |  | ihMTR | 7.337 | 0.463 | 2.630 | 0.032 | 0.020 | 0.819*** | 0.735-0.871 |
|  |  | ihMTdR1sat | 0.099 | 0.008 | 0.050 | 0.067 | 0.064 | 0.650** | 0.594-0.697 |
| CC 3 | DTI | FA | 0.404 | 0.021 | 0.092 | 0.082 | 0.028 | 0.940*** | 0.932-0.947 |
|  |  | AD (10-3) | 1.079 | 0.023 | 0.095 | 0.032 | 0.014 | 0.922*** | 0.914-0.927 |
|  |  | RD (10-3) | 0.562 | 0.023 | 0.110 | 0.047 | 0.019 | 0.911*** | 0.900-0.928 |
|  |  | MD (10-3) | 0.734 | 0.019 | 0.082 | 0.022 | 0.011 | 0.852** | 0.830-0.873 |
|  | HARDI | AFD total | 0.303 | 0.016 | 0.093 | 0.027 | 0.017 | 0.814** | 0.747-0.852 |
|  |  | NuFO | 1.119 | 0.049 | 0.334 | 0.104 | 0.050 | 0.734** | 0.716-0.749 |
|  | NODDI | ECvf | 0.433 | 0.024 | 0.109 | 0.047 | 0.030 | 0.830** | 0.797-0.856 |
|  |  | ICvf | 0.567 | 0.024 | 0.109 | 0.031 | 0.017 | 0.829** | 0.803-0.865 |
|  |  | ISOvf | 0.051 | 0.010 | 0.045 | 0.112 | 0.114 | 0.627** | 0.605-0.657 |
|  |  | OD | 0.262 | 0.013 | 0.057 | 0.131 | 0.045 | 0.944*** | 0.939-0.949 |
|  | MTI | MTR | 21.464 | 0.589 | 3.284 | 0.017 | 0.014 | 0.690** | 0.572-0.759 |
|  |  | MTsat | 3.551 | 0.151 | 1.042 | 0.058 | 0.060 | 0.555* | 0.501-0.599 |
|  |  | ihMTR | 7.596 | 0.767 | 3.905 | 0.032 | 0.019 | 0.785** | 0.617-0.863 |
|  |  | ihMTdR1sat | 0.107 | 0.012 | 0.068 | 0.071 | 0.064 | 0.639** | 0.542-0.683 |
| CST | DTI | FA | 0.480 | 0.022 | 0.110 | 0.080 | 0.036 | 0.863*** | 0.849-0.881 |
|  |  | AD (10-3) | 0.510 | 0.021 | 0.100 | 0.043 | 0.021 | 0.865*** | 0.843-0.878 |
|  |  | RD (10-3) | 1.150 | 0.029 | 0.170 | 0.056 | 0.029 | 0.879*** | 0.861-0.890 |
|  |  | MD (10-3) | 0.720 | 0.017 | 0.090 | 0.030 | 0.017 | 0.877*** | 0.856-0.895 |
|  | HARDI | AFD total | 0.330 | 0.020 | 0.120 | 0.046 | 0.028 | 0.792** | 0.759-0.835 |
|  |  | NuFO | 1.300 | 0.052 | 0.260 | 0.099 | 0.050 | 0.693** | 0.675-0.716 |
|  | NODDI | ECvf | 0.320 | 0.027 | 0.120 | 0.098 | 0.089 | 0.670** | 0.643-0.710 |
|  |  | ICvf | 0.680 | 0.027 | 0.120 | 0.036 | 0.026 | 0.680** | 0.656-0.716 |
|  |  | ISOvf | 0.100 | 0.012 | 0.060 | 0.136 | 0.125 | 0.704** | 0.685-0.738 |
|  |  | OD | 0.220 | 0.017 | 0.100 | 0.147 | 0.071 | 0.852** | 0.839-0.864 |
|  | MTI | MTR | 22.680 | 0.480 | 2.540 | 0.019 | 0.014 | 0.701** | 0.657-0.736 |
|  |  | MTsat | 3.350 | 0.136 | 0.620 | 0.055 | 0.054 | 0.533* | 0.504-0.568 |

| | | | | | | | | | |
|---|---|---|---|---|---|---|---|---|---|
| | | ihMTR | 9.660 | 0.503 | 2.650 | 0.038 | 0.024 | 0.769** | 0.724-0.843 |
| | | ihMTdR1sat | 0.120 | 0.009 | 0.040 | 0.076 | 0.060 | 0.647** | 0.579-0.699 |
| IFOF | DTI | FA | 0.390 | 0.023 | 0.150 | 0.021 | 0.030 | 0.918*** | 0.906-0.924 |
| | | AD (10-3) | 0.590 | 0.024 | 0.110 | 0.040 | 0.016 | 0.894*** | 0.881-0.904 |
| | | RD (10-3) | 1.120 | 0.032 | 0.240 | 0.115 | 0.018 | 0.895*** | 0.882-0.907 |
| | | MD (10-3) | 0.770 | 0.021 | 0.090 | 0.075 | 0.012 | 0.855** | 0.835-0.872 |
| | HARDI | AFD total | 0.270 | 0.017 | 0.090 | 0.033 | 0.019 | 0.838** | 0.803-0.867 |
| | | NuFO | 1.200 | 0.041 | 0.290 | 0.095 | 0.050 | 0.721** | 0.703-0.734 |
| | NODDI | ECvf | 0.460 | 0.023 | 0.090 | 0.032 | 0.026 | 0.824** | 0.804-0.849 |
| | | ICvf | 0.540 | 0.023 | 0.090 | 0.113 | 0.019 | 0.824** | 0.802-0.846 |
| | | ISOvf | 0.060 | 0.010 | 0.050 | 0.069 | 0.118 | 0.642** | 0.618-0.661 |
| | | OD | 0.260 | 0.019 | 0.150 | 0.033 | 0.053 | 0.918*** | 0.911-0.925 |
| | MTI | MTR | 22.140 | 0.525 | 2.430 | 0.057 | 0.013 | 0.746** | 0.688-0.775 |
| | | MTsat | 3.490 | 0.150 | 0.930 | 0.041 | 0.057 | 0.609* | 0.579-0.640 |
| | | ihMTR | 7.490 | 0.407 | 2.220 | 0.031 | 0.019 | 0.798** | 0.745-0.836 |
| | | ihMTdR1sat | 0.100 | 0.008 | 0.050 | 0.017 | 0.060 | 0.660** | 0.613-0.689 |

0.006

Descriptive and consistency statistics for microstructure measures. For AD, RD, and MD units of Mean, SD and Range is 10-3 mm2/s; CVb = averaged between-subject, CVw =

**Supplementary Table 2 Consistency metrics for MRI measurements according to fiber population.**

CVb = averaged between-subject, CVw = averaged within-subject, ICC = averaged across sessions.

| | | | ICC | | | CVw | | | CVb | | |
|---|---|---|---|---|---|---|---|---|---|---|---|
| | Model | Metrics | Average | Multi | Single | Average | Multi | Single | Average | Multi | Single |
| AF | DTI | FA | 0,86 | 0,90 | 0,90 | 0,057 | 0,039 | 0,037 | 0,106 | 0,075 | 0,081 |
| | | AD | 0,84 | 0,87 | 0,88 | 0,024 | 0,015 | 0,018 | 0,040 | 0,024 | 0,033 |
| | | RD | 0,84 | 0,88 | 0,88 | 0,026 | 0,015 | 0,020 | 0,042 | 0,027 | 0,036 |
| | | MD | 0,78 | 0,86 | 0,84 | 0,017 | 0,011 | 0,012 | 0,024 | 0,018 | 0,020 |
| | HARDI | AFD total | 0,77 | 0,86 | 0,84 | 0,030 | 0,019 | 0,022 | 0,042 | 0,033 | 0,035 |
| | | NuFo | 0,59 | 0,62 | 1,00 | 0,101 | 0,020 | 0,000 | 0,127 | 0,026 | 0,000 |
| | NODDI | ECvf | 0,74 | 0,83 | 0,81 | 0,039 | 0,027 | 0,028 | 0,044 | 0,034 | 0,038 |
| | | ICvf | 0,74 | 0,83 | 0,81 | 0,028 | 0,018 | 0,020 | 0,037 | 0,028 | 0,031 |
| | | ISOvf | 0,57 | 0,68 | 0,69 | 0,124 | 0,093 | 0,091 | 0,105 | 0,081 | 0,094 |
| | | OD | 0,87 | 0,89 | 0,91 | 0,069 | 0,037 | 0,054 | 0,124 | 0,068 | 0,103 |
| | MTI | MTR | 0,69 | 0,82 | 0,77 | 0,019 | 0,012 | 0,014 | 0,023 | 0,018 | 0,019 |
| | | MTsat | 0,77 | 0,87 | 0,83 | 0,030 | 0,019 | 0,022 | 0,044 | 0,034 | 0,037 |
| | | ihMTR | 0,47 | 0,66 | 0,60 | 0,092 | 0,058 | 0,068 | 0,071 | 0,058 | 0,063 |
| | | ihMTdR1sat | 0,57 | 0,73 | 0,68 | 0,098 | 0,062 | 0,073 | 0,093 | 0,073 | 0,080 |
| CC3 | DTI | FA | 0,90 | 0,93 | 0,91 | 0,049 | 0,032 | 0,036 | 0,109 | 0,072 | 0,089 |
| | | AD | 0,87 | 0,89 | 0,88 | 0,023 | 0,013 | 0,018 | 0,044 | 0,023 | 0,037 |
| | | RD | 0,87 | 0,91 | 0,88 | 0,028 | 0,013 | 0,024 | 0,052 | 0,027 | 0,046 |
| | | MD | 0,80 | 0,89 | 0,83 | 0,018 | 0,010 | 0,014 | 0,026 | 0,018 | 0,023 |
| | HARDI | AFD total | 0,74 | 0,85 | 0,78 | 0,026 | 0,015 | 0,021 | 0,033 | 0,023 | 0,029 |
| | | NuFo | 0,62 | 0,62 | 1,00 | 0,082 | 0,013 | 0,000 | 0,127 | 0,018 | 0,000 |
| | NODDI | ECvf | 0,78 | 0,87 | 0,81 | 0,037 | 0,022 | 0,031 | 0,049 | 0,033 | 0,044 |
| | | ICvf | 0,78 | 0,87 | 0,81 | 0,026 | 0,015 | 0,021 | 0,038 | 0,026 | 0,034 |
| | | ISOvf | 0,63 | 0,71 | 0,69 | 0,114 | 0,079 | 0,093 | 0,112 | 0,076 | 0,103 |
| | | OD | 0,90 | 0,92 | 0,92 | 0,069 | 0,030 | 0,062 | 0,145 | 0,066 | 0,125 |
| | MTI | MTR | 0,64 | 0,80 | 0,70 | 0,020 | 0,012 | 0,016 | 0,023 | 0,016 | 0,020 |
| | | MTsat | 0,75 | 0,87 | 0,79 | 0,029 | 0,016 | 0,024 | 0,043 | 0,030 | 0,038 |
| | | ihMTR | 0,42 | 0,65 | 0,52 | 0,094 | 0,056 | 0,076 | 0,072 | 0,055 | 0,067 |
| | | ihMTdR1sat | 0,54 | 0,72 | 0,62 | 0,100 | 0,060 | 0,081 | 0,094 | 0,068 | 0,085 |
| CST | DTI | FA | 0,85 | 0,84 | 0,89 | 0,042 | 0,038 | 0,028 | 0,088 | 0,062 | 0,067 |
| | | AD | 0,85 | 0,88 | 0,86 | 0,025 | 0,021 | 0,018 | 0,048 | 0,031 | 0,037 |
| | | RD | 0,86 | 0,89 | 0,87 | 0,035 | 0,024 | 0,029 | 0,062 | 0,039 | 0,051 |
| | | MD | 0,86 | 0,90 | 0,85 | 0,022 | 0,019 | 0,016 | 0,035 | 0,027 | 0,027 |
| | HARDI | AFD total | 0,77 | 0,85 | 0,80 | 0,031 | 0,025 | 0,024 | 0,047 | 0,037 | 0,038 |
| | | NuFo | 0,66 | 0,64 | 1,00 | 0,057 | 0,019 | 0,000 | 0,106 | 0,024 | 0,000 |
| | NODDI | ECvf | 0,66 | 0,75 | 0,73 | 0,089 | 0,076 | 0,066 | 0,097 | 0,081 | 0,076 |
| | | ICvf | 0,67 | 0,76 | 0,73 | 0,030 | 0,022 | 0,024 | 0,039 | 0,029 | 0,033 |
| | | ISOvf | 0,70 | 0,81 | 0,74 | 0,125 | 0,089 | 0,106 | 0,136 | 0,099 | 0,120 |
| | | OD | 0,84 | 0,83 | 0,90 | 0,082 | 0,051 | 0,069 | 0,157 | 0,080 | 0,134 |
| | MTI | MTR | 0,68 | 0,74 | 0,74 | 0,017 | 0,012 | 0,013 | 0,021 | 0,015 | 0,017 |
| | | MTsat | 0,75 | 0,79 | 0,84 | 0,027 | 0,021 | 0,021 | 0,040 | 0,030 | 0,034 |
| | | ihMTR | 0,50 | 0,61 | 0,60 | 0,062 | 0,046 | 0,050 | 0,059 | 0,043 | 0,051 |
| | | ihMTdR1sat | 0,62 | 0,66 | 0,75 | 0,068 | 0,053 | 0,054 | 0,080 | 0,057 | 0,069 |
| | | FA | 0,87 | 0,92 | 0,89 | 0,048 | 0,030 | 0,035 | 0,097 | 0,064 | 0,079 |

| | | | | | | | | | | | |
|---|---|---|---|---|---|---|---|---|---|---|---|
| IFOF | DTI | AD | 0,85 | 0,90 | 0,87 | 0,022 | 0,011 | 0,018 | 0,038 | 0,021 | 0,034 |
| | | RD | 0,85 | 0,91 | 0,87 | 0,025 | 0,012 | 0,022 | 0,043 | 0,024 | 0,039 |
| | | MD | 0,80 | 0,90 | 0,82 | 0,017 | 0,009 | 0,014 | 0,025 | 0,017 | 0,022 |
| | HARDI | AFD total | 0,78 | 0,89 | 0,82 | 0,028 | 0,015 | 0,023 | 0,040 | 0,029 | 0,036 |
| | | NuFo | 0,62 | 0,70 | 1,00 | 0,079 | 0,014 | 0,000 | 0,114 | 0,021 | 0,000 |
| | NODDI | ECvf | 0,77 | 0,89 | 0,80 | 0,032 | 0,016 | 0,027 | 0,042 | 0,028 | 0,038 |
| | | ICvf | 0,77 | 0,89 | 0,80 | 0,026 | 0,013 | 0,022 | 0,036 | 0,024 | 0,033 |
| | | ISOvf | 0,64 | 0,82 | 0,67 | 0,118 | 0,063 | 0,103 | 0,115 | 0,072 | 0,109 |
| | | OD | 0,88 | 0,90 | 0,90 | 0,069 | 0,029 | 0,061 | 0,123 | 0,056 | 0,110 |
| | MTI | MTR | 0,68 | 0,84 | 0,72 | 0,018 | 0,010 | 0,015 | 0,021 | 0,015 | 0,019 |
| | | MTsat | 0,75 | 0,87 | 0,79 | 0,028 | 0,015 | 0,023 | 0,039 | 0,027 | 0,035 |
| | | ihMTR | 0,49 | 0,74 | 0,57 | 0,086 | 0,046 | 0,071 | 0,070 | 0,053 | 0,066 |
| | | ihMTdR1sat | 0,56 | 0,77 | 0,64 | 0,092 | 0,049 | 0,075 | 0,088 | 0,064 | 0,080 |

Consistency of MRI measures according to fiber population. CVb = averaged between-subject, CVw = averaged within-subject, ICC = averaged across session.
